# Trait evolution and historical biogeography shape assemblages of annual killifish

**DOI:** 10.1101/436808

**Authors:** Andrew J. Helmstetter, Tom J. M. Van Dooren, Alexander S. T. Papadopulos, Javier Igea, Armand M. Leroi, Vincent Savolainen

**Author notes:** **Data archival location:** Genbank and Dryad.

## Abstract

Reconstructions of evolutionary and historical biogeographic processes can improve our understanding of how species assemblages developed and permit inference of ecological drivers affecting coexistence. We explore this approach in *Austrolebias*, a genus of annual fishes possessing a wide range of body sizes. Regional assemblages composed of different species with similar size distributions are found in four areas of eastern South America. Using phylogenetic trees, species distribution models and size data we show how trait evolution and historical biogeography have affected the composition of species assemblages. We extend age-range correlations to improve estimates of local historical biogeography. We find that size variation principally arose in a single area and infer that ecological interactions drove size divergence. This large-size lineage spread to two other areas. One of these assemblages was likely shaped by adaptation to a new environment, but this was not associated with additional size divergence. We found only weak evidence that environmental filtering has been important in the construction of the remaining assemblage with the smallest range of sizes. The repeated assemblage structures were the result of different evolutionary and historical processes. Our approach sheds light on how species assemblages were built when typical clustering approaches may fall short.

## INTRODUCTION

Coexistence of related species is shaped by the effects of different ecological, evolutionary and historical processes. These include speciation (Nosil 2012; Warren et al. 2014; Mittelbach and Schemske 2015) and extinction, local ecological processes such as competition (Hardin 1960; Pigot and Tobias 2012) and environmental filtering (Mouillot et al. 2007; Lebrija-Trejos et al. 2010) and processes independent of phenotype, e.g. random dispersal (Gotelli and McGill 2006). Their effects on coexistence differ across spatial and temporal scales (Webb et al. 2002). Ecological processes acting over short time scales (e.g., resource competition) can contribute to the selective pressures that drive trait evolution and ecological speciation over longer time scales (Langerhans and Riesch 2013), e.g. character displacement (Brown and Wilson 1956; Losos 1990; Schluter and McPhail 1992). Understanding the interplay between ecological and evolutionary forces is essential in order to determine the processes affecting species coexistence.

The importance of evolutionary and historical biogeographic processes in community assembly is increasingly acknowledged (Gerhold et al. 2018) but they remain relatively understudied in community ecology when compared to local and recent ecological processes (Warren et al. 2014; Mittelbach and Schemske 2015). In community phylogenetics (Webb et al. 2002), phylogenetic reconstructions are used to characterise assemblages and to predict the ecological processes at work in them. An assemblage that is phylogenetically overdispersed is usually inferred to be structured by competition, while phylogenetically clustered assemblages are thought be shaped by environmental filtering. However, a single ecological process can have variable effects on phylogenetic relatedness and trait variation in an assemblage (Cavender-Bares et al. 2009). Typical community phylogenetic approaches based on phylogenetic relatedness are unable to discriminate between processes effectively as they implicitly assume simple evolutionary models (Weber et al. 2017). They are also susceptible to interpreting historical effects on current geographic distributions as evidence for ecological and evolutionary processes (Warren et al. 2014).

Trait evolution is an essential component in the formation of species assemblages (Webb et al. 2002, Cavender-Bares et al. 2004; Kraft et al. 2007). However, when models of trait evolution are applied, they are often relatively simple: convergence in traits is often not modelled while conserved traits are modeled using a Brownian motion (BM) model of evolution (e.g. Kraft et al. 2007). Tools that permit inference of interactions between ecological and evolutionary processes are still in development (Weber et al. 2017), but it is possible to investigate variability of evolutionary processes across a phylogenetic tree without having to resort to heuristic sampling algorithms (Kraft et al. 2007). Methods can be used to detect whether species traits in an assemblage are attracted to more than a single phenotypic optimum, by identifying selection regime shifts (Butler and King 2004). They can also be used to infer whether traits of different species converge towards the same optimum (Ingram and Mahler 2013; Oke et al. 2017; Speed and Arbuckle 2017). Taking advantage of these approaches is important because trait convergence can also be the result of independent, random divergence from distant starting points (Webb et al. 2002; Stayton 2008). By separating the history of selection regimes from evolutionary random walks we can improve our understanding of the driving forces behind trait evolution and coexistence (Oke et al. 2017).

Historical biogeography is also intrinsic to how species assemblages form but is often neglected in empirical studies (Warren et al. 2014; Mittelbach & Schemske 2015). For example speciation in sympatry or parapatry (i.e. non-allopatric) produces co-occurring sister species while speciation in allopatry does not, although this patterns will change over time due to post-speciation range shifts. The most common way biogeography is included in phylogenetic studies is through ancestral range estimation (ARE) models, albeit at a coarse level. Species are assigned to predefined areas, which are used to estimate different types of ‘cladogenetic events’ that imply varying levels of geographic proximity during divergence. For example, ‘founder-event speciation’ (long-distance dispersal followed by isolation and speciation; Matzke et al. 2014) lies on one end of the geographic continuum while ‘within area speciation’ is as close as these models can get to the other end. Identifying speciation in sympatry is particularly difficult (Papadopulos et al. 2011; Igea et al. 2015; Martin et al. 2015) and requires the examination of evidence beyond the scope of phylogenetic comparative methods (Coyne and Orr 2000). Nevertheless, ARE can identify instances where species ranges may have overlapped during divergence, which can be useful to develop further hypotheses. We can attempt to refine phylogenetic predictions of geographic overlap during divergence by using methods that make use of species range maps such as the age range correlation approach (ARC; Barraclough and Vogler 2000). Making use of ARC alongside ARE can help us obtain a more complete picture of the biogeographic context and the processes that shaped species assemblages.

To better understand how trait evolution and historical biogeography have shaped species assemblages we need a study group with some natural replication, i.e., multiple assemblages with comparable traits but different sets of related species. One such group is a genus of freshwater fishes, *Austrolebias* (Costa 1998; 2006). These annual killifishes live in seasonal freshwater pools and wetlands on the South American grasslands and floodplains. During the reproductive season, a mating pair will dive into the muddy substrate of the pond, oviposit and fertilise their eggs. As the dry season commences, ponds dry and the adults die. Eggs persist within the soil, going through a single or several stages of diapause until hatching is triggered by the wet season rains (Wourms 1972). Once hatched, fish grow rapidly to adulthood. The annual life-cycle of *Austrolebias* restricts them to a specific habitat where interactions are typically between congenerics. This is useful because their close relatedness may allow us to link history, ecology and evolution more easily.

*Austrolebias* are principally found in the seasonal ponds distributed throughout basins of the La Plata and Paraguay Rivers, the Negro River drainage of the Uruguay river basin and the Patos-Merin lagoon drainage system (Fig. 1). These areas are referred to as their areas of endemism (Costa 2010) and we consider assemblages as the species that are found in each of these areas. The ranges of *Austrolebias* species can be small or large within each area of endemism and each range can be characterised by environmental variables. These distinguish species by what we could call their realised environmental niche. We refer to quantitative representations of species differences in these variables simply as environmental niches. The life-cycle of *Austrolebias* relies heavily on the seasonal environment and the extent to which their environmental niche can vary and have changed in the past is unknown. It is possible that this has influenced the formation of current assemblages and the trait variation within. We will apply a commonly used heuristic and study changes in environmental niche using the same methods as for regular traits (Münkemüller et al. 2015), while we are aware that these abstract traits are not defined at the organismal level and can change without evolution.

**Figure 1.**
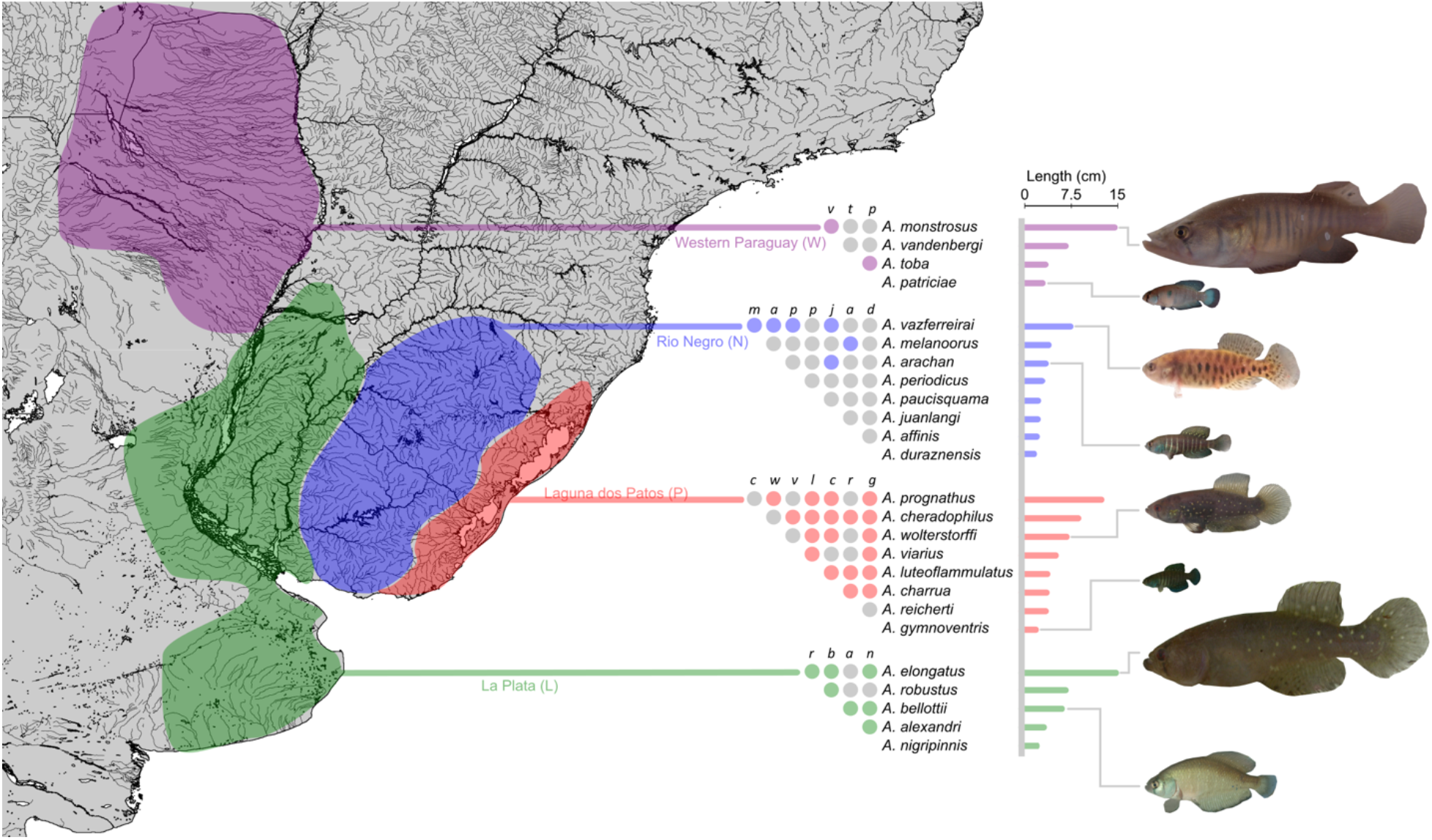
A map depicting the four main areas of endemism in *Austrolebias*, inferred by drawing shapes around sampling locales for species in each region. The most northern region, in purple, is the area west of the River Paraguay (W). The basin of the La Plata river and its delta are in green (L). The drainage of the Negro River, part of the River Uruguay basin is represented in blue (N). The Patos-Merin Lagoon region is highlighted in red (P). The species used in this study are shown on the right side of this figure, grouped by area. *Austrolebias apaii* and *A. cinereus* are not shown. Body size measurements are depicted as bars alongside species names. Local patterns of species co-occurrence are shown in the pairwise table to the left of species names. Coloured dots indicate that a record of a pond with both species was found. Images of fishes are approximately to scale. All fish images are of males except *A. vazferreirai*.

Within each area of endemism species vary substantially in adult body size, resulting in an approximate replication of trait distributions (Fig. 1). The largest species can grow to more than 150mm in length while the smallest can be less than 25mm when mature (Costa 2006) and differently-sized species are known to locally coexist in each area (Fig. 1). Variation in body size is particularly interesting because size influences resource use in freshwater systems (Woodward and Hildrew 2002) and stronger competition is expected between those species with similar body sizes because they use the same resources (MacArthur and Levins 1967; May and MacArthur 1972). Diet has already been shown to be linked to body size in *Austrolebias* (Laufer et al. 2009; Arim et al. 2010; Ortiz and Arim 2016) and the largest species are known to prey upon the smaller species (Costa 2009) indicating that differently sized species occupy distinct niches in a pond. An ecological study within a single area found that local community structure was determined more by body size than species identity (Canavero et al. 2014). A reconstruction of body size evolution is fundamental for understanding how *Austrolebias* assemblages were built.

We use models of trait evolution and historical biogeography to disentangle the processes that led to the replicated body size distributions across *Austrolebias* assemblages. When we detect a selection regime shift for the optimal size of a species relative to its ancestor, models of historical biogeography and a decision tree can help determine the process that has generated this change (Fig. 2). We suggest three types of scenario that can be detected when using this approach. Scenario (A1) occurs when biogeographic analyses at the regional and local scales predict that a particular speciation event was allopatric. Concurring regime shifts for size and environmental niche following divergence will indicate that size evolution is linked to a substantial change in the environmental niche. Therefore, the size change can be interpreted as an adaptation to a new environment. Scenario (B1) is similar to (A1) but without the shift in environmental niche. Here a number of processes (e.g., dispersal syndromes (Stevens et al. 2014); accelerated evolution upon colonisation (Aubret 2015)) are compatible with the scenario (B1) so we leave the inferred process open. Scenarios (A2) and (B2) are identical to (A1) and (B1) but account for the possibility that speciation events classified as within-area at the regional scale may be revealed as allopatric at the local scale. Scenario (C) involves a selection regime shift for size and a shift for environmental niche while speciation is non-allopatric. Competition and adaptation to a new environment can contribute to this scenario however their presence is not required such that we cannot reliably infer either process. Scenario (D) occurs when speciation is non-allopatric. A regime shift for body size alone may indicate that trait divergence was driven by divergent or disruptive selection towards different size optima through ecological interactions (Fig. 2).

**Figure 2.**
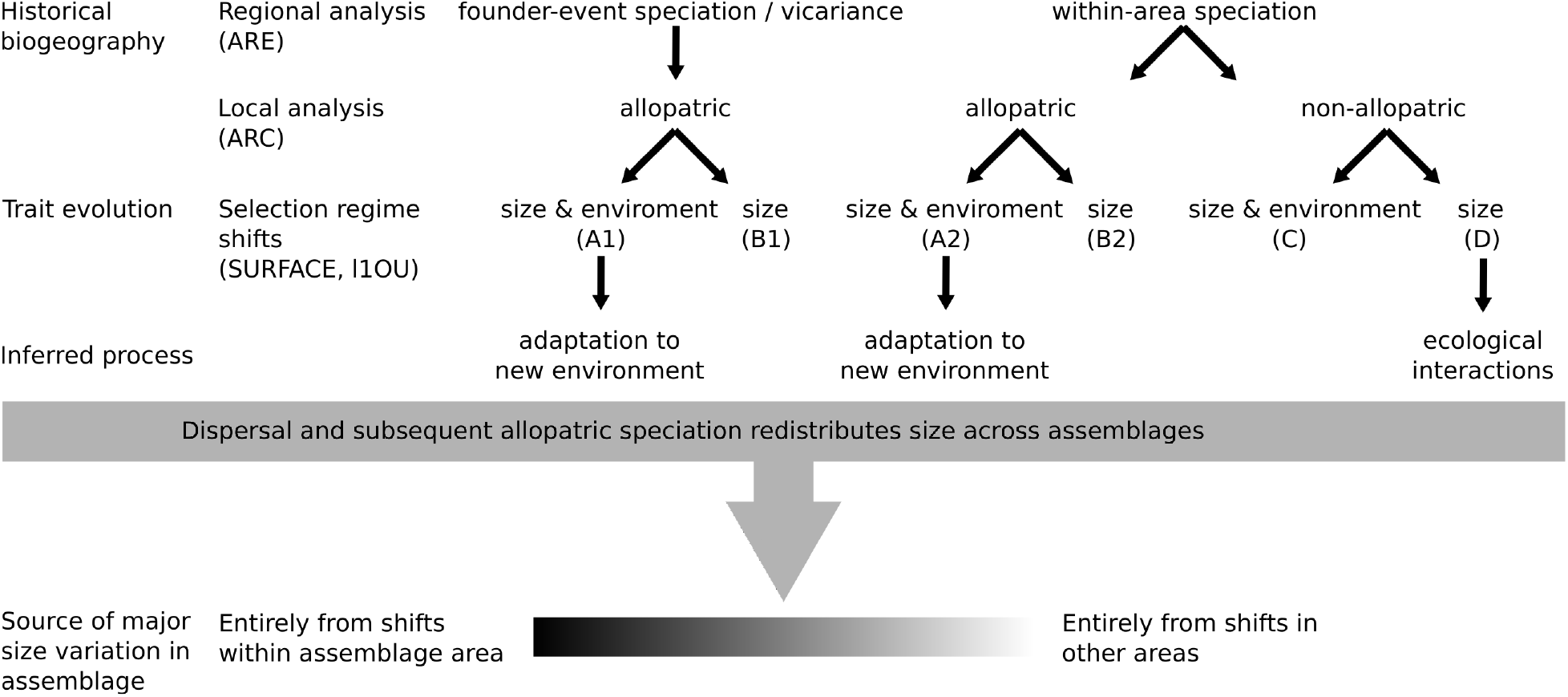
A decision tree showing how we can combine historical biogeography and trait evolution with selection regime shifts to understand how replicated size distributions across assemblages were generated. We examine the historical biogeography at each cladogenetic event in the tree. Initially we do this at the regional level, classifying divergence events based on results from BioGeoBEARS analysis (ARE). We subsequently use ancestral range correlations (ARC) to see if this finer-scale approach can improve upon ARE. When a selection regime shift in size or a shift in environmental niche (“environment” in the diagram) is inferred just after divergence, we can determine whether the pattern corresponds to adaptation to a new environment or to ecological interactions (e.g. competition). Several patterns of shifts do not allow clear inference of processes. Once we have established how major size variation generated by selection regime shifts has arisen we can then examine if and how it has been redistributed throughout the assemblages.

Identifying the occurrence of these scenarios can shed light on how trait variation arose but may not fully explain how similar trait distributions were composed in each assemblage. We can again use historical biogeographic approaches to understand how size variation generated in one assemblage became distributed over others. For example, size-diverged species that originated through competition (D) can emigrate from their original assemblage to occupy the same ecological niche in a different assemblage and speciate in allopatry. This would preempt the scope for scenarios (C) or (D) in the colonised assemblage (Rueffler et al. 2006). We can rank assemblages according the contribution of dispersal to size variation (Fig. 2). At one extreme all major size variation in an assemblage came from shifts within or upon colonisation of the assemblage area. At the opposite extreme major size variation appeared exclusively by dispersal.

Here we sequence nuclear (nDNA) and mitochondrial DNA (mtDNA) markers in 26 species to reconstruct the evolutionary history of *Austrolebias* and address four main questions. In conjunction with new phylogenetic trees we perform traditional community phylogenetics to ask whether we find phylogenetic clustering or overdispersion in the assemblages. We collate size and occurrence data to model species distributions, estimate environmental niches and identify shifts in selection optima to ask whether selection regime shifts in body size and shifts in environmental niche occurred. We reconstruct regional biogeographic events and complement this with an extension of the age-range correlation (Barraclough and Vogler 2000) to test if we can make predictions of local historical biogeography. Finally, when selection regime shifts are detected, we can attempt to uncover how size variation arose, determine the sources of size variation per assemblage (Fig. 2) and answer our last question: Have the similar size patterns across *Austrolebias* assemblages been generated in the same way? We found that although trait distributions are similar, the major processes that shaped size variation in each assemblage were not.

## MATERIAL AND METHODS

### DNA sequencing

We used the Qiagen DNeasy Blood & Tissue kit to extract genomic DNA from 60 individuals of 26 *Austrolebias* species obtained from field sampling and populations maintained at CEREEP in Nemours, France. For each extraction, 15mg of fin tissue was used, except for small fish where we complemented with muscle tissue. All individuals were sequenced on an ABI 3130xL Genetic Analyzer for fragments of ectoderm-neural cortex protein 1 gene (*enc1*), recombination activating gene 1 (*rag1*), SH3 and PX domain containing 3 (*snx33*), rhodopsin (*rh1*), and three fragments of 28S ribosomal DNA (28SrRNA). Three mitochondrial genes were sequenced: 12S ribosomal DNA (12S-rRNA), 16S ribosomal DNA (16S-rRNA) and cytochrome b (*cytB*). Primers for the amplification and sequencing of three fragments of 28S-rRNA and the mitochondrial genes 12S-rRNA, 16S-rRNA and *cytB* were taken from Van Dooren et al. (2018). For the enc1 gene primers from (Li et al. 2007) were used. We designed new primers for *snx33*, *rag1* and *rh1* using Primer3 (Untergasser et al. 2012; Table S1).

To improve taxonomic coverage, our dataset for tree inference was supplemented with sequence data from Van Dooren et al. (2018). When sequences from different sources were combined, the sequences were of same source population (Table S2). Additional sequences for *Pterolebias longipinnis* (GenBank accession EF455709, KC702007, KC702072) and *Hypsolebias magnificus* (KC701989, KC702056, KC702122) were downloaded from GenBank to be used as outgroups. Sequences were aligned in Geneious (v6.1, Biomatters) using the MAFFT alignment plugin (MAFFT v7.017; Katoh et al. 2002) and low quality ends were trimmed. The combined nuclear matrix is composed of 65 individuals and 3,128 aligned characters (*enc1*: 706, *rag1*: 651, *snx33*: 543, 28S-rRNA: 724, *rh1*: 504). The combined mitochondrial matrix comprises 65 individuals and 1,563 characters (*cytB*: 720, 12S-rRNA: 345, 16S-rRNA: 498).

### Phylogenetic inference

Sequences representing 63 *Austrolebias* individuals and two outgroups were used to build nDNA and mtDNA-based trees as well as a tree constructed using nDNA and assuming a multispecies coalescent with *BEAST. Phylogenetic tree inference was conducted using BEAST2 v2.4.3 (Bouckaert et al. 2014). We initially used a model testing approach to choose the appropriate substitution model for each partition but BEAST2 analyses failed to converge. Therefore, we used the relatively simple substitution model HKY+G as suggested in Drummond and Bouckaert (2015) and empirical base frequencies for each gene/partition. For mtDNA and nDNA trees we used linked trees with separate substitution models for each locus and a single uncorrelated relaxed clock model. The exception were the 28S rRNA fragments, which were run with linked substitution models across fragments. In our *BEAST analyses for the coalescent, substitution, clock and tree models were unlinked and the 28s fragments were run as a concatenated alignment. Yule tree priors were used for all analyses.

Phylogenetic trees were calibrated using divergence times of *Austrolebias* with two outgroups and normal priors; *Pterolebias longpinnis* (mean = 46.79 million years ago (Ma); s.d. = 3.45) and *Hypsolebias magnificus* (mean = 16.54 Ma; s.d. = 2.75). Both secondary calibrations were taken from Helmstetter et al. (2016). We constrained clades containing (i) *Austrolebias* and *Hypsolebias* and (ii) only *Austrolebias* to be monophyletic to match the topology in (Helmstetter et al. 2016). BEAST2 and *BEAST were run for 5.0 × 10^7^ generations and trees were sampled every 5,000 generations. We repeated runs three times for each dataset. Tracer v1.6 (Drummond and Rambaut 2007) was used to identify at which point stationarity had been reached and LogCombiner (Bouckaert et al. 2014) to combine trees from separate runs. The relevant percentage of trees was discarded as burn-in to ensure that effective sample size was greater than 200 for all parameters. Finally, TreeAnnotator (Bouckaert et al. 2014) was used to generate the maximum clade credibility (MCC) tree. For downstream analyses we converted mtDNA and nDNA trees to species trees with the GLASS algorithm (Liu et al. 2009) using the speciesTree function in R library ‘ape’ (Paradis et al. 2004). This created a pair of species trees we could use along the *BEAST species tree for further analyses.

### Phylogenetic clustering

We tested for the presence of phylogenetic clustering or overdispersion in the assemblages of each of the four areas of endemism (detailed below). We used mean pairwise phylogenetic distance (MPD, Webb et al. 2002) and simulated null distributions for this statistic per assemblage (R library picante; Kembel et al. 2010).

### Location data, species ranges & species distribution modelling

We aggregated coordinates of ponds where *Austrolebias* have been observed using primary publications, the Global Biodiversity Information Facility (https://www.gbif.org/), information on amateur egg trading websites and data shared by hobbyist collectors or from our own collection trips (Table S3, updated until March 2017). Using Google Earth, we obtained coordinates for all locations, identified duplicates and verified pond locations when images of their locations were available. At each location in the dataset, we examined species co-occurrence and added matrices in Figure 1 showing which species pairs co-occur in our data complemented with reports in Loureiro et al. (2015).

We characterised species ranges in two ways. First, we used the four areas of endemism that distinguish the assemblages in our analysis (Costa 2010). These are: the western region of the Paraguay River basin (Western Paraguay or W), the lower La Plata River basin and the middle-lower Uruguay River basin (La Plata or L), the Negro River drainage of the Uruguay River basin and the upper/middle parts of the Jacuí, Santa Maria, Jahuarão and Quaraí river drainages (Negro River or N) and the Patos-Merin lagoon system including the southern coastal plains (Patos Lagoon or P) (Fig. 1). A fifth area of endemism is found the middle section of the Iguaçu River basin. It is home to a single species, *A. carvalhoi,* for which we do not have data, so it was not included.

Second, we built species distribution models (SDMs) using the above data and MaxEnt (version 3.3.3k; Phillips et al. 2006; Phillips and Dudík 2008) to more accurately determine species ranges. We used the current global climate data layers of 19 bioclimatic variables and altitude taken from WorldClim (www.worldclim.org) at a spatial resolution of 30 arcseconds (approximately 1 km^2^), 10 variables characterising soil composition and two variables with river basin characteristics (Table S4). As most locations where *Austrolebias* have been sampled are at roadsides, we sampled background points from a raster file of roads at the same resolution (CIESIN 2013). We ran a PCA on the unstandardised environmental variables and used the principal component scores (PC) as covariates in MaxEnt. This produced ranges with fewer spurious predicted occurrences at a large distance from the known capture sites than PC from standardised environmental variables. Range sizes were calculated by applying a threshold to the logistic output of MaxEnt, at which the sum of the sensitivity (true positive rate) and specificity (true negative rate) is highest. When a cell possessed a habitat suitability score above this threshold, the species was counted as present in that cell.

### Environmental niches

We characterised the environmental niche of each species with statistics of the environmental variables listed above. We followed an approach similar to the outlying mean index (OMI) analysis proposed by Dolédec et al. (2000). First, environmental variables were all standardised to zero mean and standard deviation equal to one. Then their values at the known locations per species were averaged. We weighed locations by presence/absence as local abundances are unknown and the different ecological roles in the genus might affect abundances independent of environmental preferences (e.g. large piscivore species are less numerous in ponds). On the resulting values a PCA was carried out which did not account for phylogenetic relatedness among species as we did not want to impose an evolutionary model *a priori*.

### Body size data

We represented species size by a measure for asymptotic length. Growth is indeterminate in fish and age information is often not available with size measurements. Body size data were obtained by taking the largest known field measurements of adult male standard length (SL) for each species from the literature and our own field records (full dataset in Table S5).

### Prediction of ancestral ranges

Ancestral range estimation (ARE) was conducted using the R package ‘BioGeoBEARS’ (Matzke 2014). The biogeographic events this approach assesses are within-area speciation, within-area subset speciation, range expansion, range contraction, vicariance and founder event speciation (summarised in Dupin et al. 2016). Some events occur within an area defining an assemblage, others imply a dispersal or vicariance event. ‘BioGeoBEARS’ fits models similar to the Dispersal-Extinction-Cladogenesis (DEC) model (Ree and Smith 2008), the BayArea (Bayesian Inference of Historical Biogeography for Discrete Areas) model (Landis et al. 2013) and the Dispersal-Vicarance (DIVA) model (Ronquist 1997), with and without founder-event speciation (indicated by +j). We ran separate sets of analyses where the maximum number of areas composing ancestral ranges was restricted to two (maximum observed) or four (maximum possible). We performed model selection based on Akaike Information Criterion corrected for sample size (AICc), to identify which models best fit the data. We ran analyses where ranges were restricted to include only adjacent areas. For example P could only share a range with N, while N could be part of a range with both L and P (Fig. 1). Finally, Costa (2010) suggested that marine transgressions during the middle to late Miocene may have connected W, N and P, allowing for dispersal among these areas. We therefore ran a separate set of models that classified these areas as adjacent between 15 and 11 Ma to determine whether including this transgression led to a better fit. We implemented Biogeographical Stochastic Mapping (BSM; Dupin et al. 2016) to predict the frequencies of different biogeographic events, using the best models for each tree (500 simulations).

### Selection regime shifts for size and environmental niche shifts

We used SURFACE (Ingram and Mahler 2013) to identify convergent evolution in body size and convergent environmental niche shifts. This program fits Ornstein–Uhlenbeck (OU) models with one or several optima and applies adapted Information Criterion comparisons in a stepwise manner to identify regime shifts on a phylogenetic tree and determines whether these tend towards the same optima. We limited our analysis of the species environmental niche to the first two PC of environmental variables. We also ran l1OU (Khabbazian et al. 2016) to see whether an alternative method of identifying evolutionary shifts corroborated our SURFACE results.

### Ancestral range correlations

Inferring historical biogeography using areas of endemism and ARE can oversimplify spatial patterning of the actual species ranges and their overlaps. We used the age-range correlation (ARC) approach of Barraclough and Vogler (2000) which can handle detailed range data. Simulations have shown that it has some power to discriminate geographic modes of speciation (Barraclough and Vogler 2000). ARCs are regression models of range overlap on phylogenetic node age. Descendant ranges are pooled to calculate range overlap per node and overlap is expected to increase (decrease) with node age for nodes that originated by allopatric (sympatric) speciation. A major fault in the current inference of the probabilities of different modes is that these are inferred from the estimated intercept of the regression across pooled data. However, there are many combinations of geographical speciation modes that can produce the same intercept. We used mixture regressions in the ‘flexmix’ R package (Grün and Leisch 2008) to analyse age-range overlap data. Nodes of a phylogenetic tree can be assigned to different clusters of events representing different geographical speciation modes and with the intercept and node age slope estimated per cluster. Mixtures fitted consisted of 1-3 component regression models.

Range overlaps were calculated as fractional overlap of the smallest range size following Barraclough and Vogler (2000). For one species, *A. paucisquama*, we were unable to infer a reasonable SDM so we could not include this species in our ARC analyses. After an exploration of logit and other models, we analysed the data as mixtures of gaussian random variables with linear regressions. These produced fewer spurious results and an absence of strong effects of parameterisation details. We constrained the residual variances of all mixture components to be equal. We otherwise obtained regressions with very small error variances, which were not observed in process simulations (Barraclough and Vogler 2000). We inferred range overlap intercepts by bootstrapping the preferred mixture model, which was selected on the basis of comparing ICL (Integrated Completed Likelihood criterion) between models.

## RESULTS

### Phylogenetic inference

The topologies of nDNA (Fig. 3A) and mtDNA (Fig. 3C) trees were well supported with posterior probabilities (PP) > 0.9 in 74% & 69% of nodes, respectively. Support in the coalescent tree (Fig. 3B) was markedly lower (38% of nodes with PP > 0.9, 58% of nodes with PP > 0.75). The topology of our mtDNA tree is similar to two other recent mtDNA trees (García et al. 2014; Van Dooren et al. 2018). Estimates of the age of the genus overlapped between the trees. It was calculated to be 18.37 Ma (95% HPD = 14.03, 22.94) in the nDNA tree, slightly older in the mtDNA tree at 19.50 Ma (95% HPD = 14.63, 23.53) and 12.48 Ma (95% HPD = 8.47, 16.61) for the coalescent tree. These estimates are similar to a study of the Cynolebiini (Costa et al. 2017) which dated the crown age of *Austrolebias* at approximately 10-15 Ma. García et al. (2014) did not use secondary calibrations and estimated a more recent crown age of approximately 8 Ma.

**Figure 3.**
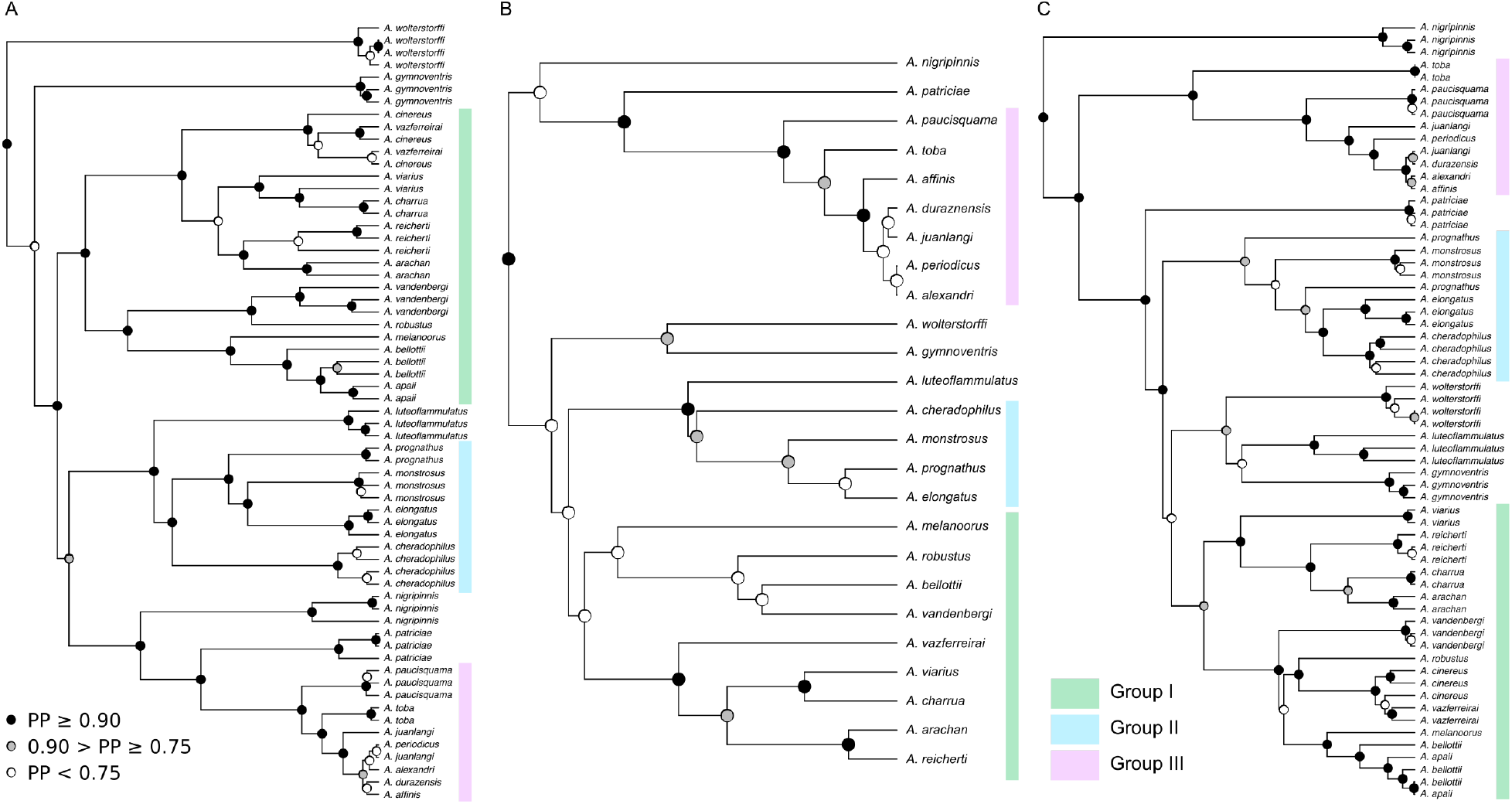
Maximum clade credibility trees from Bayesian analyses of (A) nDNA, (B) coalescent nDNA and (C) mtDNA. Black circles indicate a posterior probability (PP) from 0.90 – 1.00 and grey circles indicate a PP from 0.75 to 0.90. PP < 0.75 is depicted as a white node. Highlighted, colour-coded sections represent three major clades that are recovered in all trees. Relationships within clades may vary between trees. Branches units are in millions of years.

We consistently recovered three major groups within *Austrolebias* in all trees (Fig. 3), which did not include all species. Several species were non-monophyletic in our nDNA and mtDNA trees including *A. apaii, A. bellottii, A. cinereus, A. juanlangi* & *A. viarius*. *Austrolebias bellottii* and *A. apaii* were treated as the same species because *A. apaii* is a junior synonym of *A. bellottii* (García et al. 2012). In some cases *A. cinereus* individuals were more closely related to *A. vazferreirai* than to their conspecifics. These species are also morphologically similar (Costa 2006) and *A. cinereus* is known from only a single population. We decided to merge *A. vazferreirai* and *A. cinereus* for the comparative analyses in this paper including the *BEAST analysis. Our mtDNA topology differed extensively from our nrDNA tree (Fig. 3), which had substantial effects on downstream inference. There are many reasons why the mtDNA history may not accurately reflect the species tree (Shaw 2002; Ballard and Whitlock 2004; see discussion). Similarly, the poor support in the coalescent tree means that it is likely unreliable for inference. Therefore, while we performed each analysis using all three trees, we focus on summarising the results based on the nDNA tree.

### Phylogenetic clustering

We found significant phylogenetic clustering for the Negro River assemblage (MPD score 22.5, null expectation 28.4, *z* = −2.50, *p* = 0.02). Classically, this would be taken as evidence that environmental filtering was the major process involved in building that assemblage. For the other assemblages, there is no significant phylogenetic clustering or overdispersion.

### Ancestral range estimation

The best-fitting ARE models were DIVA+j for all trees. The parameters and likelihoods of each model are summarised in Table S6. These models revealed that jump-dispersal of lineages between areas was common (Fig. 4, Fig. S1). Accordingly, biogeographic stochastic mapping found frequencies in the range of 47-60% of events (Fig. S2). Likelihood ratio tests revealed that models with the jump-dispersal parameter conferred significantly higher likelihood to the data than those without (Table S7) indicating that founder-event speciation was important in shaping the distribution of *Austrolebias* species. One striking observation was that the most northern area of endemism, Western Paraguay (W), was colonised by *Austrolebias* in three (coalescent & mtDNA) (Fig. S1) or four (nDNA) independent instances (Fig. 4).

**Figure 4.**
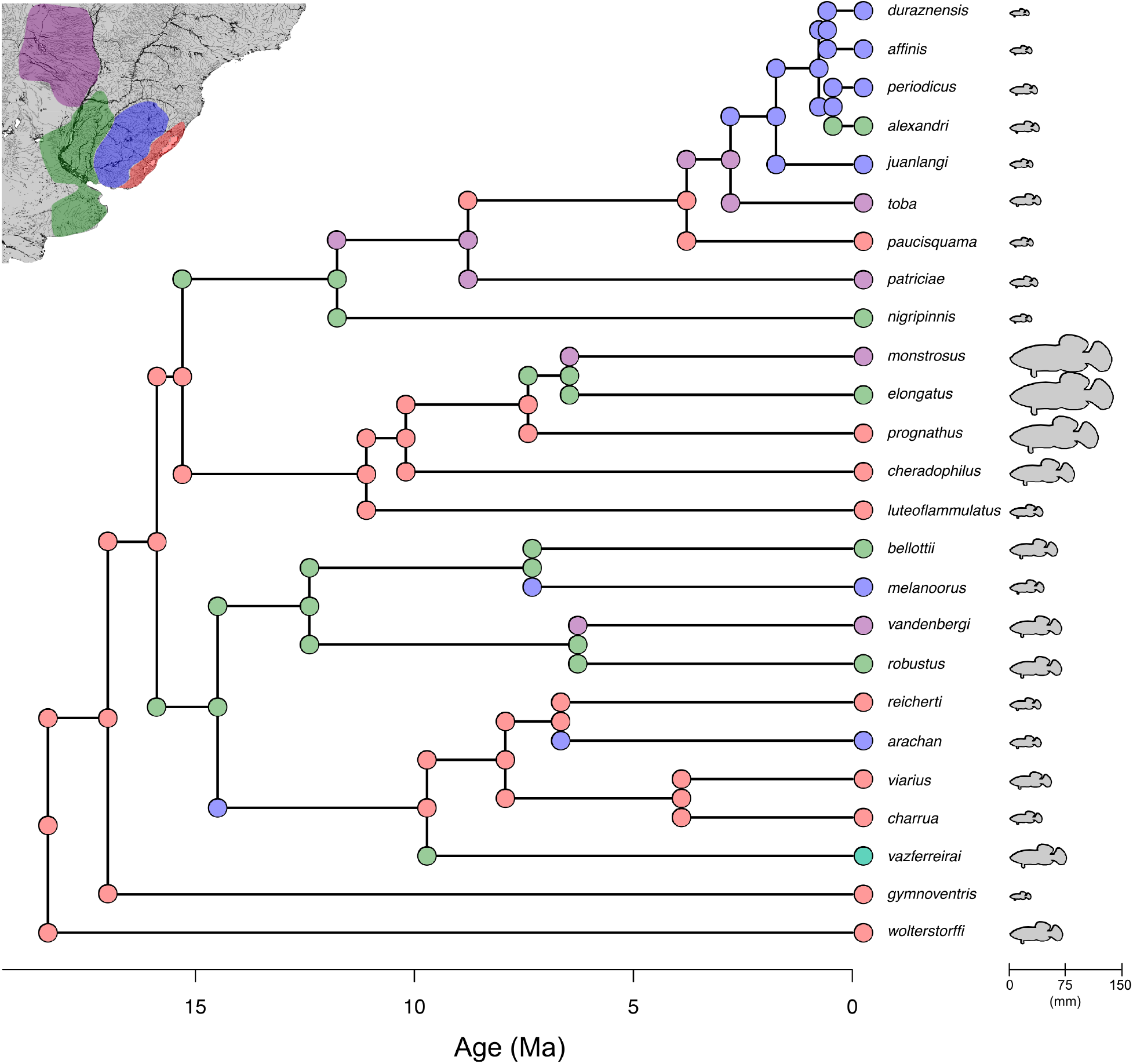
A biogeographic estimation of ancestral ranges using the trimmed nDNA tree and the best fitting model (DIVA+j with a maximum of two areas) in BioGeoBEARS. Most probable states are shown on each node and corner of the tree, current ranges are shown on the tips. Colours correspond to regions as depicted in the map in the top left and Figure 1. The current range of *A. vazferreirai* consists of two areas - Negro River and La Plata (coloured as teal). The body lengths of each species are shown as silhouettes to scale to the right of tree.

Speciation was also estimated within areas of endemism. Expected numbers of within-area speciation events were particularly high in Patos Lagoon (an average of 4.4 events per simulation) and Negro River (2.7) while the other areas had averages below one. Vicariance events were rare, probably due to the rarity of ancestral ranges with more than one area. Within-area subset speciation is not included in the DIVA model. Neither adjacency matrices nor changing the maximum number of areas significantly improved the likelihood (< 2 difference in AICc) when compared to the simplest set of models (Table S6).

### Size evolution

Our ancestral state reconstructions (Fig. 5A) show substantial divergence in size. The four largest *Austrolebias* species form a single clade. Models consistently found a shift in selection optimum towards this clade (Group II in Fig. 3) in all trees, just after the divergence of the smaller *A. luteoflammulatus*. (Fig. 5A, Fig. S3, AICc decrease relative to single optimum model mtDNA:13.3, nDNA: 13.6 coalescent: 11.1). An additional shift towards an intermediate optimum was inferred in the mtDNA tree (AICc decrease 0.9), in a clade containing *A. bellottii, A. melanoorus*, *A. robustus, A. vandenbergi* and *A. vazferreirai* (Fig. S3C). The remaining species in each tree tended towards another, smaller optimum and were not subjected to a shifted selection optimum for size throughout their history. Model comparisons using l1OU supported a shift in the clade with the largest species. In agreement with the pattern in AICc values observed, it found no evidence of any further shift (Fig. S4). Therefore selection regime shifts with convergence of optimal sizes have not occurred.

**Figure 5.**
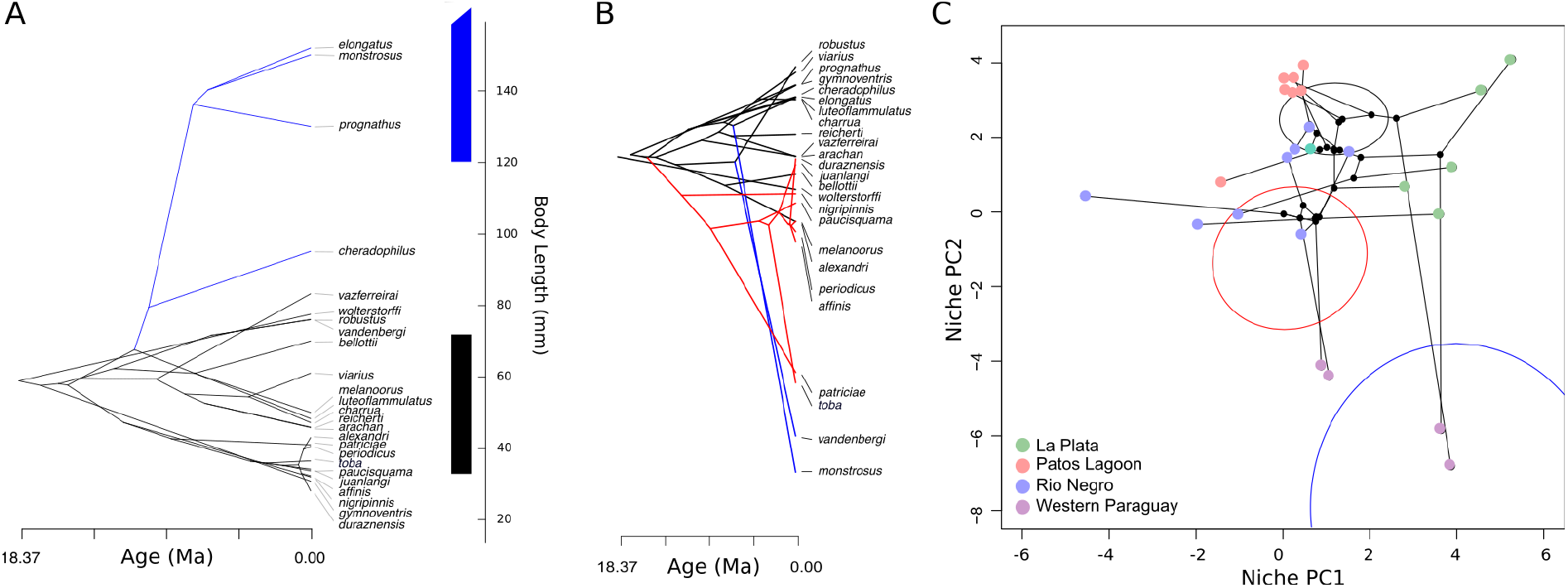
(A) Ancestral state reconstruction of body size across *Austrolebias* using stabletraits (Elliot and Mooers 2014). Ancestral values shown are the medians of the posterior distributions of size. Colours correspond to the selective optima species are tending towards as inferred by SURFACE. Bars on the right show the 95% confidence intervals of these optima. The interval for the larger size (blue) is truncated. (B) The nDNA tree arranged by the species values for principal component two (PC2) of the environmental niche. Coloured branches show the clades in which niche shifts have taken place. (C) A scatterplot of the species averages of the first two principal components. Points are tips and nodes of the nDNA tree, connected by branches representing the inferred topology. Tip values are coloured based on current areas of endemism as shown in Figure 1. Black points represent median ancestral values reconstructed from the bivariate species averages using stabletraits. 95% confidence ellipses for the combination of environmental niche values characterising each regime estimated by SURFACE are shown in their respective colours.

### Environmental niches

The first two PCs of the environmental niche variables explained 60% of species variation. We found that species cluster to some degree by area of endemism (Fig. 5B, C), and along two axes. Patos Lagoon species and Western Paraguay occupy two extremes for PC2, and all other species and ancestors group along an axis mostly determined by PC1. We assessed the scores of the two PCs. PC1 consists largely of soil information and temperature seasonality, plus information on the driest and coldest season. Western Paraguay and La Plata are similar for this PC and have less acidic soil with more silt, a larger seasonal range in temperatures and less annual precipitation. PC2 contains information relating to average temperatures, precipitation seasonality and precipitation/temperature during the wettest month and the warmest season. This axis primarily distinguishes Western Paraguay from the other areas, with warmer temperatures, increased precipitation seasonality and more rain in the warmest season.

We assessed whether there were shifts and convergent changes among the environmental niche changes. SURFACE analyses using PC1 & PC2 revealed two regime shifts (Fig 5, Fig. S3; AICc differences nDNA: 12.5 mtDNA: 11.8 coalescent: 17.0). All four species from Western Paraguay were subject to shifts associated with their dispersal into the area (Panels B & C in Fig. 5). According to the models based on the nDNA tree, convergent shifts towards a new environmental niche occurred independently when *A. vandenbergi* and *A. monstrosus* colonised Western Paraguay. A group containing *A. patriciae* and *A. toba* tended towards a second different environmental niche at a less decreased value of PC2 (Fig. 5B, C). Four out of seven species in the Negro assemblage converge to that same niche regime. l1OU analyses inferred just two shifts, for *A. monstrosus* and *A. vandenbergi* (Fig. S5). None of the regime shifts for environmental niche are associated with selection regime shifts for size, therefore scenarios (A1) and (A2) where size evolves due to adaptation to a new environment (Fig. 2) did not occur.

### Age-range correlations

Ranges varied considerably in size from the very large ones of *A. bellottii* and *A. nigripinnis* to the extremely small of *A. toba* and *A. affinis* (Fig. S6). In general, the area under the curve (AUC) values for the training data were above 0.95. Among mixture models fitted to range overlaps, models were preferred with node age effects (Table S8 contains ICL values of fitted models). For the nDNA tree (Fig. 6), a mixture with three components representing different modes of speciation was preferred. The intercepts of these three components are −0.02 (s.e. 0.03), 0.24 (s.e. 0.04) and 0.59 (s.e. 0.09). The first value corresponds to allopatric speciation, the other two to varying degrees of non-allopatric speciation. For the coalescent tree, a mixture with three components had lowest ICL, which was only slightly smaller than the ICL of a single-component mixture. We therefore infer for the coalescent and mtDNA tree (Fig. S7) a model with a single mixture component, each time with an intercept not significantly different from zero (coalescent 0.16 (s.e. 0.11), mtDNA 0.071 (s.e. 0.101)).

**Figure 6.**
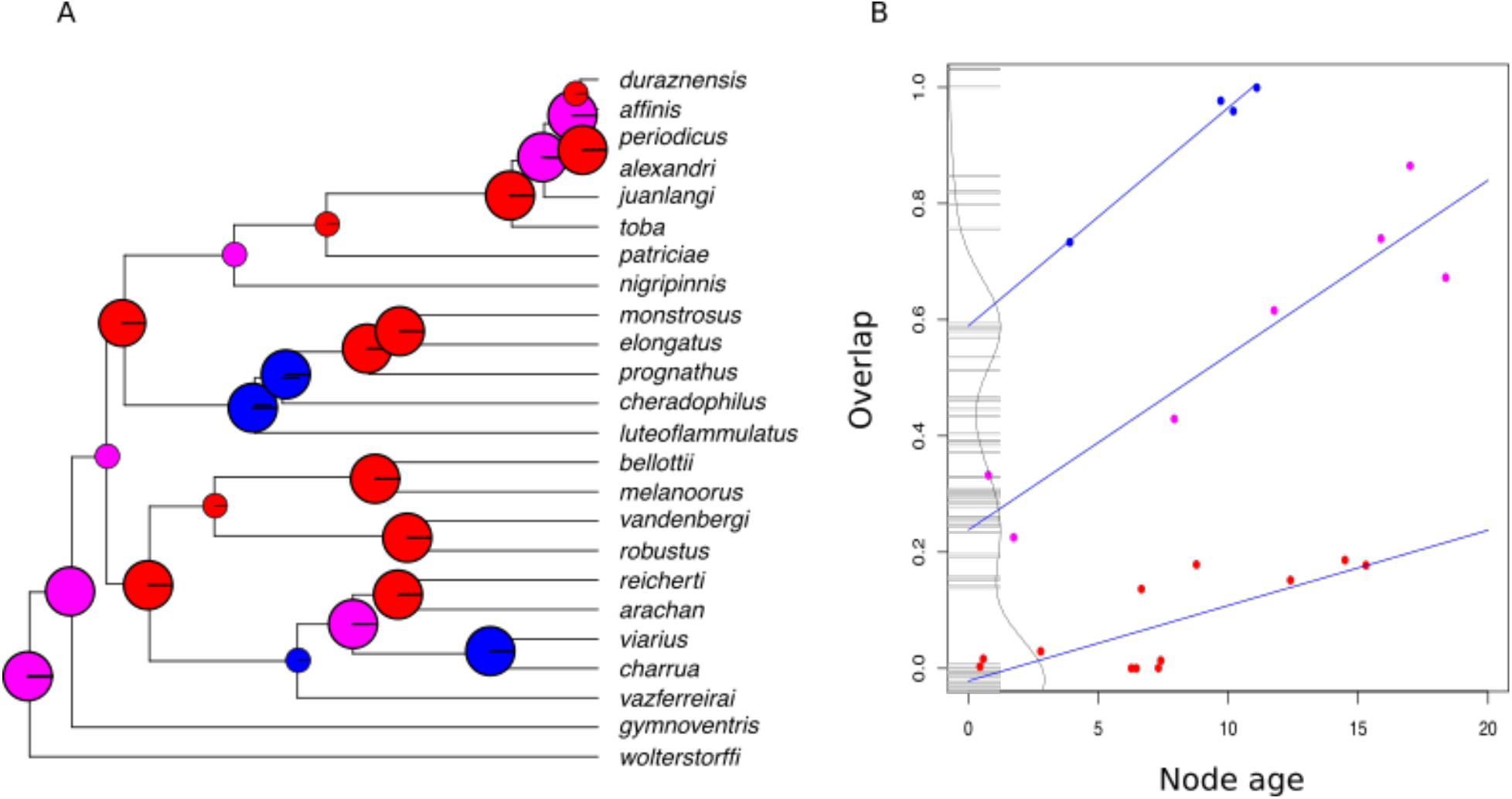
Assessments of patterns of past range overlap at cladogenetic events. (A) The nDNA tree with pie charts per node representing probabilities of assignment to mixture components. A model with three components was favoured. The component with the smallest value of overlap at the intercept is labelled red (allopatric), with increasing levels of overlap magenta and blue respectively (both non-allopatric). Nodes where we found concurrent results in our ARC and BioGeoBEARS analyses are plotted larger. (B) Scatterplot of range overlap per node on node age. Range overlap is estimated as in Barraclough and Vogler (2000) and fitted regression lines are shown for each cluster. A rug and density plot of bootstrapped intercept values is added at node age zero.

When we compare the regional and more local analyses (Fig. 4, 6A), 17 of 23 nodes showed agreement between analyses. There are three events where the regional analysis inferred nonallopatric speciation, while ARC did not. For three allopatric speciation events at the regional scale, ARC inferred non-allopatry. This combination is an impossible geography, showing limitations of historical biogeographic approaches. Most importantly, the selection regime shift for size occurred at a node where non-allopatric speciation is inferred. We can conclude that ecological interactions (C) drove this size divergence.

## DISCUSSION

### A new phylogenetic hypothesis for *Austrolebias*

The trees presented in this study are the most comprehensive inferences in *Austrolebias* to date. Our nDNA tree is the first multi-locus tree based on nuclear DNA for this genus and provides novel insight into species relationships. We find major differences between nDNA and mtDNA trees. Incomplete lineage sorting and introgression are possible causes of incongruence among different molecular markers and the maternal inheritance of mtDNA may lead to a different evolutionary history (Maddison 1997; Funk and Omland 2003; Ballard and Whitlock 2004). As an example we take *A. luteoflammulatus*, a species with different yet strongly supported placements in nDNA and mtDNA trees. It is most closely related to *Austrolebias gymnoventris* and *A. wolterstorffi* in the mtDNA tree, two species it co-occurs with. Conversely, it is sister to the clade of the largest species in the nDNA and coalescent trees. This might be evidence of mitochondrial capture - where the mtDNA of a species is completely replaced by that of another and known to occur in fish (Chan and Levin 2005; Willis et al. 2013). Other cases of discordance can be explained by low posterior probabilities in one or more phylogenetic trees. We found lower support in the coalescent tree when compared to the others and difficulty in resolving non-monophyletic species such as *A. viarius* and *A. juanlangi* (Fig. 3A) may have contributed to this.

We used sequence data and/or geographic information from 26 *Austrolebias* species, while more than 40 have been described (Costa 2006; García et al. 2014). Many species not included were discovered recently and have few known sites of occurrence. Some may be junior synonyms like *A. apaii* (García et al. 2012). Furthermore, it is not possible to build accurate species distribution models with very rare species, as evidenced by *A. paucisquama*. Nevertheless, our data covered variation among *Austrolebias* species well: all extant species in the La Plata and Western Paraguay regions were included, as well as all of the largest species (Costa 2006).

### Evidence for phylogenetic clustering in one assemblage

Assessing our assemblages using a traditional community phylogenetic approach revealed that only one of the four areas deviated from null expectations. The Negro River assemblage was significantly more clustered than expected, which is usually interpreted as evidence for environmental filtering. Environmental niches of species in Negro River did not cluster strongly (Fig. 5C) and we found no robust evidence for regime shifts, perhaps because their environmental niches were similar to species in Patos Lagoon and La Plata in most cases. Alternatively, we can explain the clustering by a relatively recent diversification of several species within the assemblage and limited scope for jump-dispersal within that time frame (Warren et al. 2014). Lack of significant results in the other three areas left the processes shaping them undefined. This justifies taking a closer look at evolutionary and historical processes to try to understand how the observed size patterns came to be.

### A single selection regime shift for body size

Body size was a priori expected to be an important factor shaping species assemblages due to the similar variation within each area of endemism (Fig. 1) and the strong size effects found in Patos Lagoon communities (Canavero et al. 2014). Our selection regime analyses consistently recovered a single shift towards the largest species. We found that this shift occurred in Patos Lagoon just after *A. luteoflammulatus* (Fig. 4, 5A) diverged. *Austrolebias luteoflammulatus* is a typical small species*, A. cheradophilus* is a large omnivore and *A. prognathus* is a very large piscivore. We speculate that a change in diet (see Laufer et al. 2009; Arim et al. 2010; Ortiz and Arim 2016) may have been associated with the regime shift for size and that predation by a piscivore facilitates coexistence of competitors. In the La Plata & Patos Lagoon areas small, similar-sized *Austrolebias* species co-occur (Fig. 1), which is unexpected if competition among *Austrolebias* would be the main factor structuring communities there, so other important processes are likely at play as well. Furthermore, in Western Paraguay *Austrolebias* are found in ponds together with other annual killifishes such as *Trigonectes aplocheiloides*, *Papiliolebias bitteri* and *Neofundulus paraguayensis* (Alonso et al. 2016) and with *Cynopoecilus melanotaenia* (Canavero et al. 2014) in Patos Lagoon. These other species may have played a role in how species assemblages are constructed in some areas e.g. by preventing the establishment of additional small *Austrolebias* species in Western Paraguay.

### Convergent shifts for environmental niche

The seasonal life cycle of *Austrolebias* is strongly climate and habitat-dependent. We did not expect to find substantial variation nor shifts in environmental niche among species. In the African annual killifish genus *Nothobranchius*, only moderate environmental niche differences were found among five species (Reichard et al. 2017). In *Nothobranchius*, altitude was found to be the most important factor explaining the distribution of different clades and species whereas in *Austrolebias* environmental niche differences between species were predominantly due to differences in soil characteristics, temperature and precipitation.

Contrary to our expectations we found that environmental niches of species from Western Paraguay were markedly differentiated from the others. SURFACE analyses revealed that the colonisation of Western Paraguay coincided with substantial and convergent shifts towards new environmental niches. These seem instrumental in how the Western Paraguay assemblage was constructed. It should be investigated whether evolutionary changes are associated with them. Western Paraguay is in the far north and west of the range of *Austrolebias* species and another distinguishing characteristic seems to be that ponds fill in winter (Schalk et al. 2016), not in summer as is typical for the other assemblages (Errea and Danulat 2001). We found two shifts of different magnitudes for species inhabiting Western Paraguay, but the less extreme found by SURFACE (Fig. 5) was not recovered in l1OU analyses (Fig. S5). While the less extreme shift seems plausible, SURFACE is known to overestimate the number of shifts (Ho and Ané 2014). Observed shifts for environmental niche were not associated with selection regime shifts for size.

### Improved prediction of local historical biogeography

To improve the prediction of historical biogeography we have extended a method developed to determine the primary geographical mode of speciation in a clade (Barraclough and Vogler 2000). We used mixture regression models to cluster data into groups of nodes fitted by a shared regression. This allowed us to estimate the intercepts of the regression lines fitted to each cluster, which approximate the range overlap at cladogenesis per group. We used the assignment of nodes to mixture components to assign speciation modes to each divergence event, without actually reconstructing ancestral ranges and without pre-defined discrete biogeographic areas.

The ARC approach has been criticised in the past (Fitzpatrick and Turelli 2006) and these criticisms still apply to our extended methodology. The power of the original method is low and the discriminatory power between alternative modes of speciation disappears at older nodes in a phylogenetic tree (Barraclough and Vogler 2000). A heuristic node weighting method has been proposed (Fitzpatrick and Turelli 2006) but in trials we did, these issues seemed aggravated by the suggested modifications. Despite these caveats we have shown that ARC can be useful for examining historical biogeography at a finer scale than ARE approaches. It is worth noting that we had to evaluate properties of several model specifications to find inference results that seemed adequate. This calls for extensive simulation studies to produce guidelines for application of the approach in different taxonomic groups.

While patterns in *Austrolebias* revealed an appreciable fraction of non-allopatric speciation events, this did not reflect the pattern in *Nothobranchius* where speciation is thought to be exclusively allopatric and triggered by geological events (Dorn et al. 2014; Bartáková et al. 2015). Our ARC results corroborated our ARE analyses (Fig. 6) for most of the speciation events in the tree. Results suggest that by using ARC we could improve upon estimates of geographic proximity of incipient species during divergence. For example, the divergence of *A. duraznensis* and *A. affinis* was estimated as within-area speciation using ARE (Fig. 4) but in allopatry using ARC (Fig. 6). Additional simulations would be useful to understand factors affecting error rates of the approach, as we found three cases where ARE and ARC predicted incompatible biogeographic events.

### Repeated patterns are the result of different processes

In Figure S8 we provide a visual summary of our reconstructions per assemblage. We observed a single selection regime shift in size, indicating that there was a single source of major size variation in the genus. Historical biogeographic approaches estimated that the speciation event that preceded this shift was within-area speciation at the regional level and non-allopatric speciation at the local level, taking place in the Patos Lagoon area. We therefore predict that scenario (C) took place - size divergence was generated through ecological interactions while incipient species were in contact. It is difficult to determine exactly what kind of interaction was behind size divergence; competition is a first possibility and Van Dooren et al. (2018) suggested that cannibalism may have led to the emergence of large predator species. The species involved in this shift, (*A. cheradophilus, A. luteoflammulatus* & *A. prognathus*) are currently known to coexist within the same ponds (Fig. 1), which lends support to the idea that they coexisted during divergence. Estimating historical patterns of gene flow among these species could investigate this further.

Our best fitting ARE model (Fig. 4) reveals two major events that were key in distributing size variation across assemblages. First, a jump-dispersal event from Patos Lagoon to La Plata led to *A. elongatus,* followed by an event from La Plata to Western Paraguay which led to *A. monstrosus*. Therefore Patos Lagoon represents one extreme of size redistribution by dispersal (Fig. 2) because major size variation originated within, and Western Paraguay and La Plata another since major size variation originated in other areas. Conspicuously absent is the Negro River assemblage, which harbours no size variation generated by a selection regime shift. *Austrolebias vazferreirai* is the largest species in this assemblage and is thought to be a generalist (Costa 2006) rather than a piscivore. Negro River is the youngest assemblage (colonised a maximum of ~7.3 Ma ago compared to ~11.8 Ma in the next youngest), so there may not have been enough time for a large predator to colonise or arise.

### Perspective

Understanding the processes that lead to current species assemblages is central in the study of ecology and evolution (Hutchinson 1959), and our study adds to increasing knowledge of the importance of species interactions (Cornell and Lawton 1992) and size differences (Simberloff and Boecklen 1981). We find similarities between our results and those from other study groups. Leitao et al. (2016) showed how rare species had a disproportionate role in the functional structure of assemblages. This may also be the case in *Austrolebias,* where large species are typically rarer than smaller species (Lanés and Maltchik 2010; Lanés et al. 2014) and occupy different functional niches, therefore widening the functional richness of assemblages. Dispersal has not led to monopolisation (whereby the first colonist gains an advantage over the later immigrants by adapting to local conditions; De Meester et al 2016) because different species probably have different ecological roles due to size and functional niche divergence. Analyses of the co-occurrence of related species of hummingbirds highlighted the importance of multiple mechanisms including competition, dispersal and the environment, in shaping assemblages (Weinstein et al. 2017). Although the methodological approaches were different, we also found that these mechanisms, as well as historical biogeography, were important in generating assemblages of *Austrolebias*.

If we had taken only a traditional community phylogenetic approach we would have dismissed Patos Lagoon as uninteresting because of no significant phylogenetic clustering. Instead our approach has revealed evidence for ecological interactions that were key to the way in which *Austrolebias* assemblages develop. Using models of trait evolution beyond Brownian motion and assessing historical biogeography at multiple levels allowed us to identify that, despite the apparent similarity of *Austrolebias* assemblages, each was generated under a different process. Having characterised each area of endemism, we can use our results to make predictions on how ponds in different areas are structured.

Body size was found to be important in structuring pond assemblages in Patos Lagoon (Canavero et al. 2014), motivating our study. In Western Paraguay and La Plata, large piscivores are also present and we would expect results similar to what was found in Patos. However, the two niche regime shifts in Western Paraguay (Fig. 5) make us predict that coexistence there is structured more by environment than in Patos. Species with more extreme environmental niches such as *A. monstrosus* and *A. vandenbergi* will occupy different ponds. We predict that body size will be less important for structuring pond assemblages in the Negro River area. Inherently, there is then more scope for weak environmental filtering effects or neutral coexistence. Confirmation of our predictions would require careful assessment of co-occurrence across these areas.

## CONCLUSION

Despite their apparent similarities, *Austrolebias* assemblage compositions and histories vary greatly. Our analysis demonstrates that reconstructed selection regimes and historical biogeography can inform on ecological processes operating in assemblages. A single selection regime shift towards large body size was critical for generating the similar trait distributions observed in *Austrolebias* assemblages. It occurred during a series of nonallopatric speciation events in a single assemblage. Historical biogeographic reconstructions revealed that dispersal and subsequent allopatric speciation redistributed major size variation to other areas. Our approach can be used in other studies examining the processes composing trait-structured assemblages. We support calls for a wider application of a historical and evolutionary view on species assemblages.

## SUPPLEMENTARY INFORMATION

**Figure S1.**
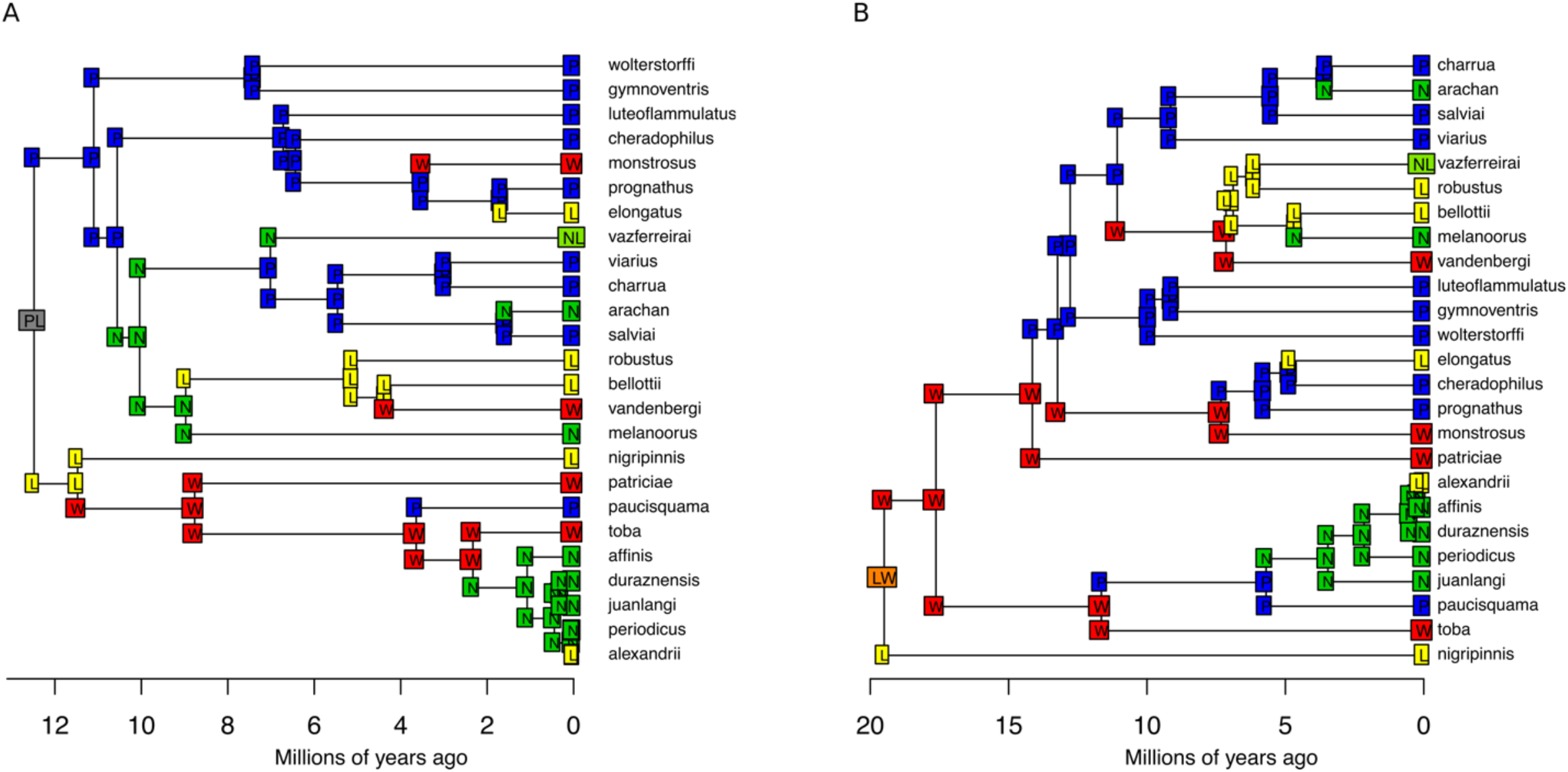
Results from ancestral range estimation using BioGeoBEARS. Most probable states under the best-fitting model for each tree are depicted on nodes with letters indicating areas of endemism. This model was DIVA+j with a maximum of two areas for the coalescent tree (A) and for the mtDNA tree (B).

**Figure S2.**
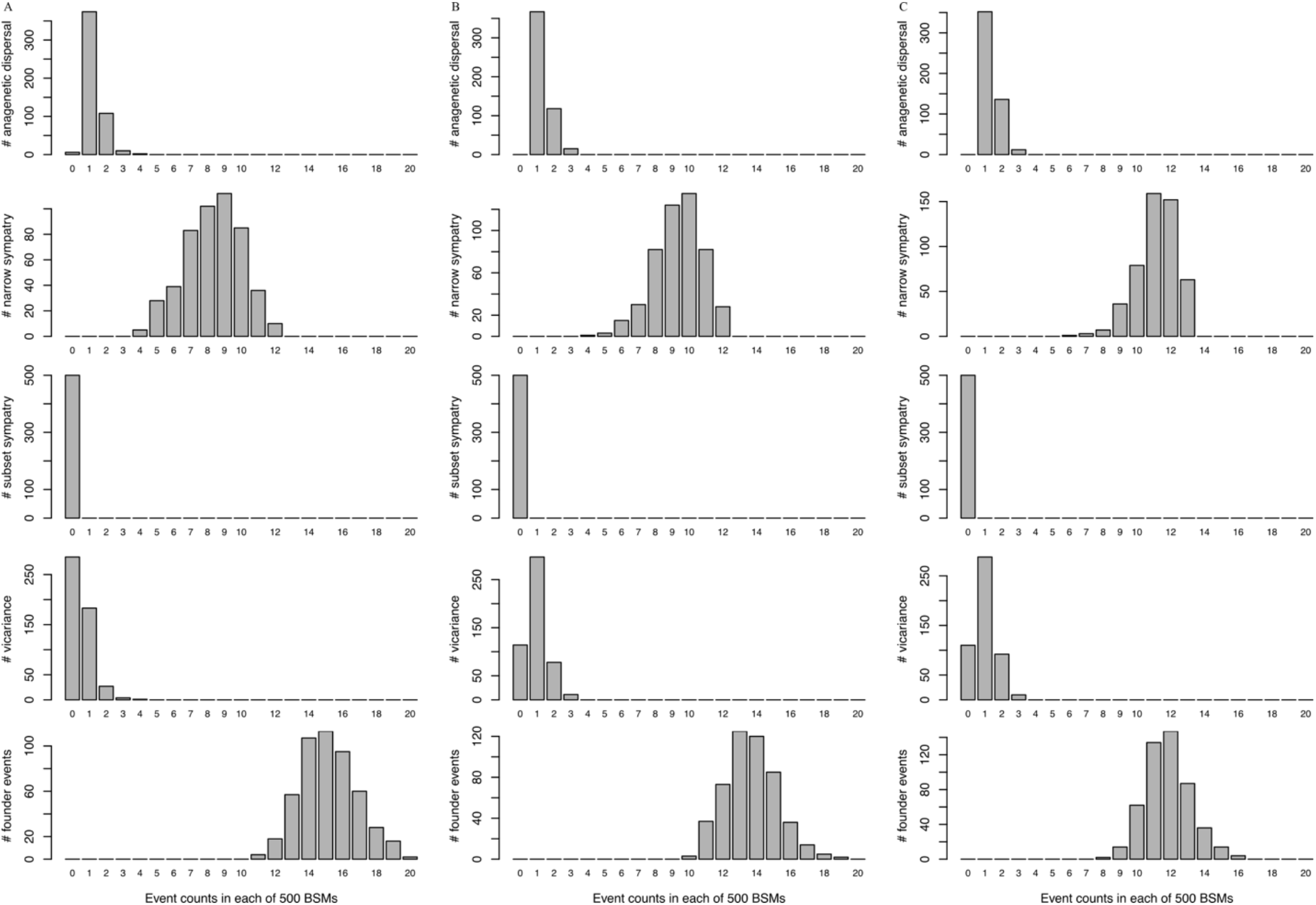
Results from 500 biogeographic stochastic mapping simulations showing frequency histograms of event counts for each event type. Results are shown for the (A) nDNA tree, (B) coalescent tree and (C) mtDNA tree.

**Figure S3.**
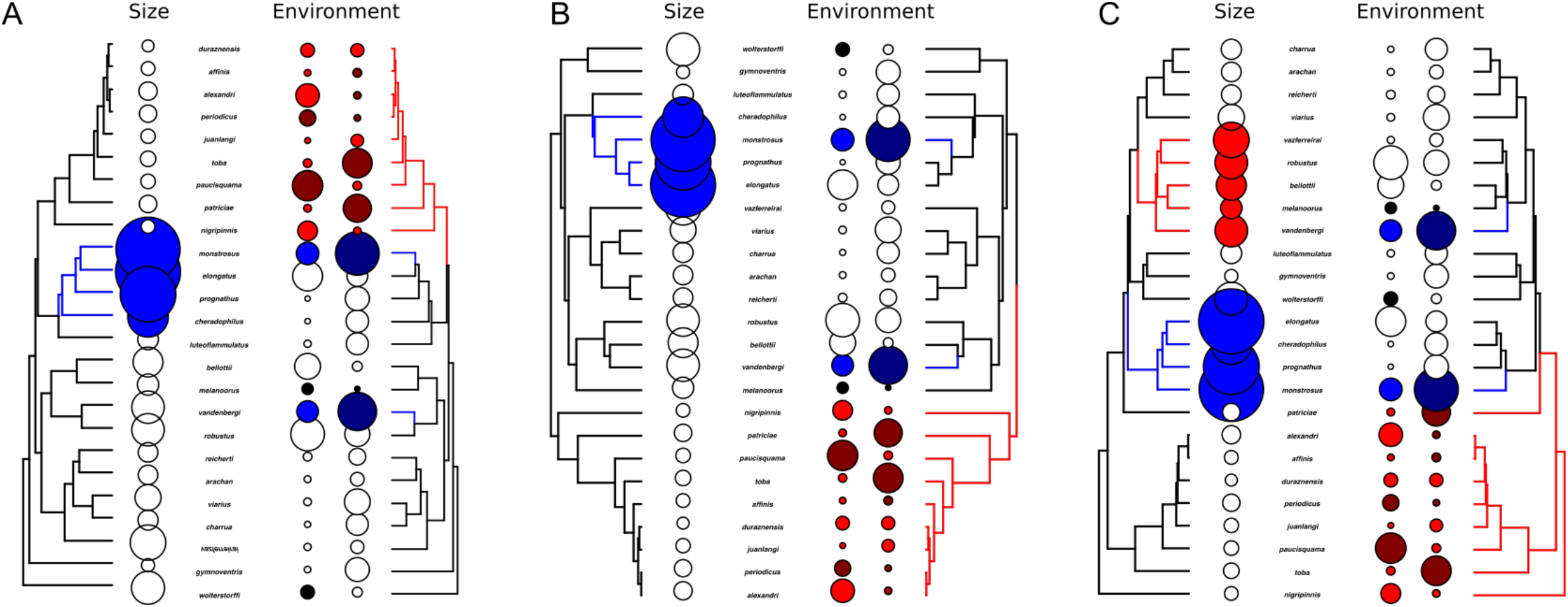
SURFACE analyses showing shifts in selective optima and trait values for body size and two environmental niche PCs using the (A) nDNA tree (B) coalescent tree and (C) mtDNA tree. Black branches represent branches where the trait is attracted to the first selective optimum for each trait, blue branches the second and red branches the third, if detected. Sizes of circles next to tips indicate magnitudes of trait values, with dark colours indicating negative magnitudes and light colours positive values. Colours correspond to relevant shifts traits are involved in.

**Figure S4.**
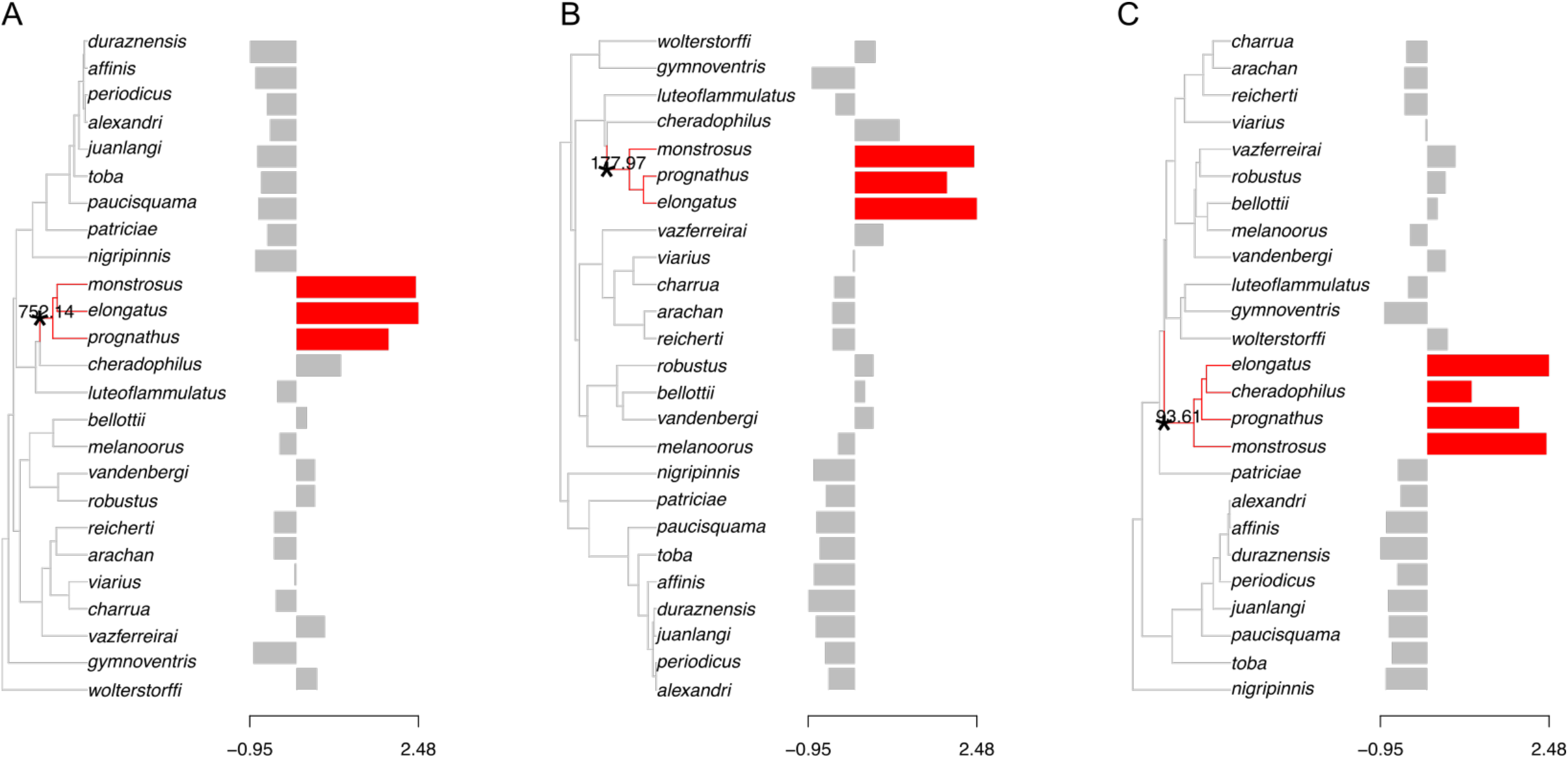
L1OU results for body size for nDNA (A), coalescent (B) and mtDNA (C) tree, where shifts are indicated by an asterisk and a change in branch colour. Shift magnitudes are shown at the tree edge of the shift. The sample data are shown in the bar chart to the right of each tree, with the data normalized.

**Figure S5.**
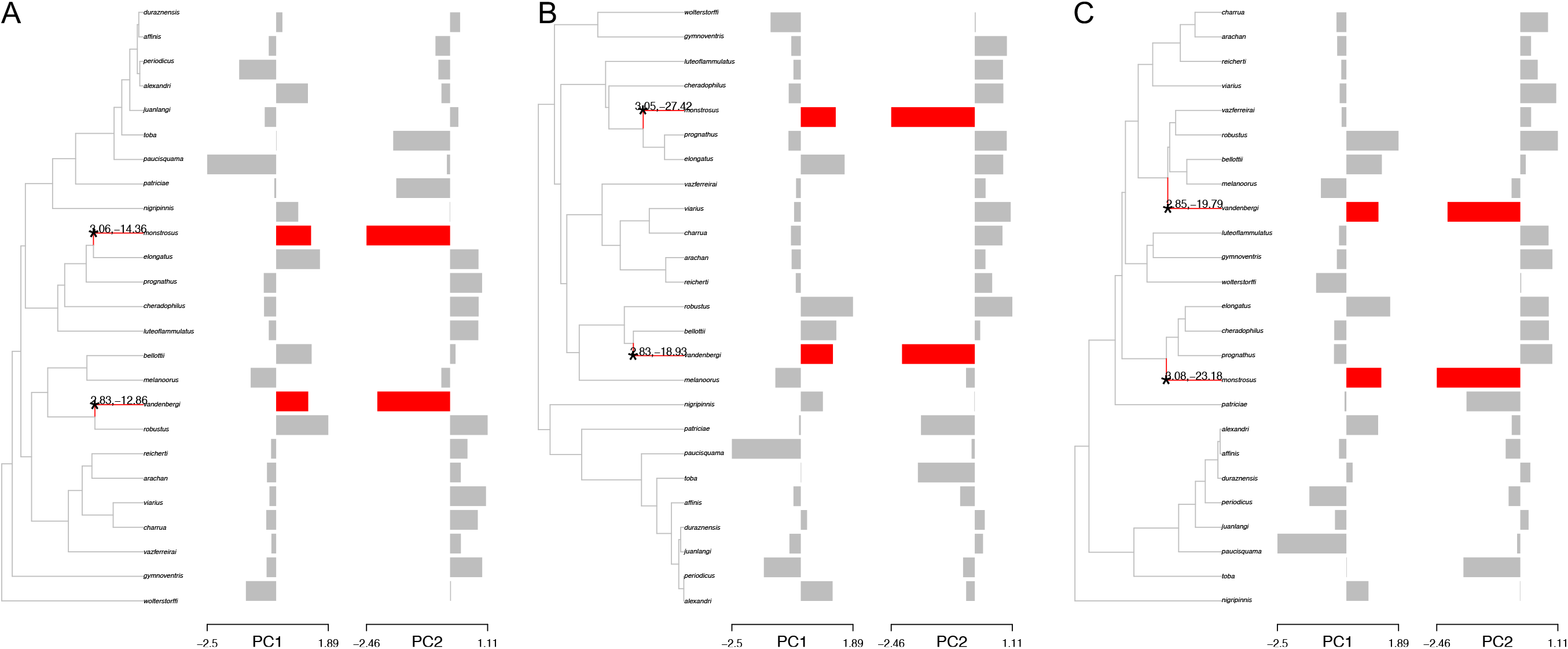
L1OU results for two environmental niche PCs for nDNA (A), coalescent (B) and mtDNA (C) tree, where shifts are shown by an asterisk and affected tree branches by a change in branch colour. The shift magnitudes are shown at the edge of the shift. The sample data for each PC are shown in the two bar charts to the right of each tree. The niche traits were normalized for these bar charts.

**Figure S6.**
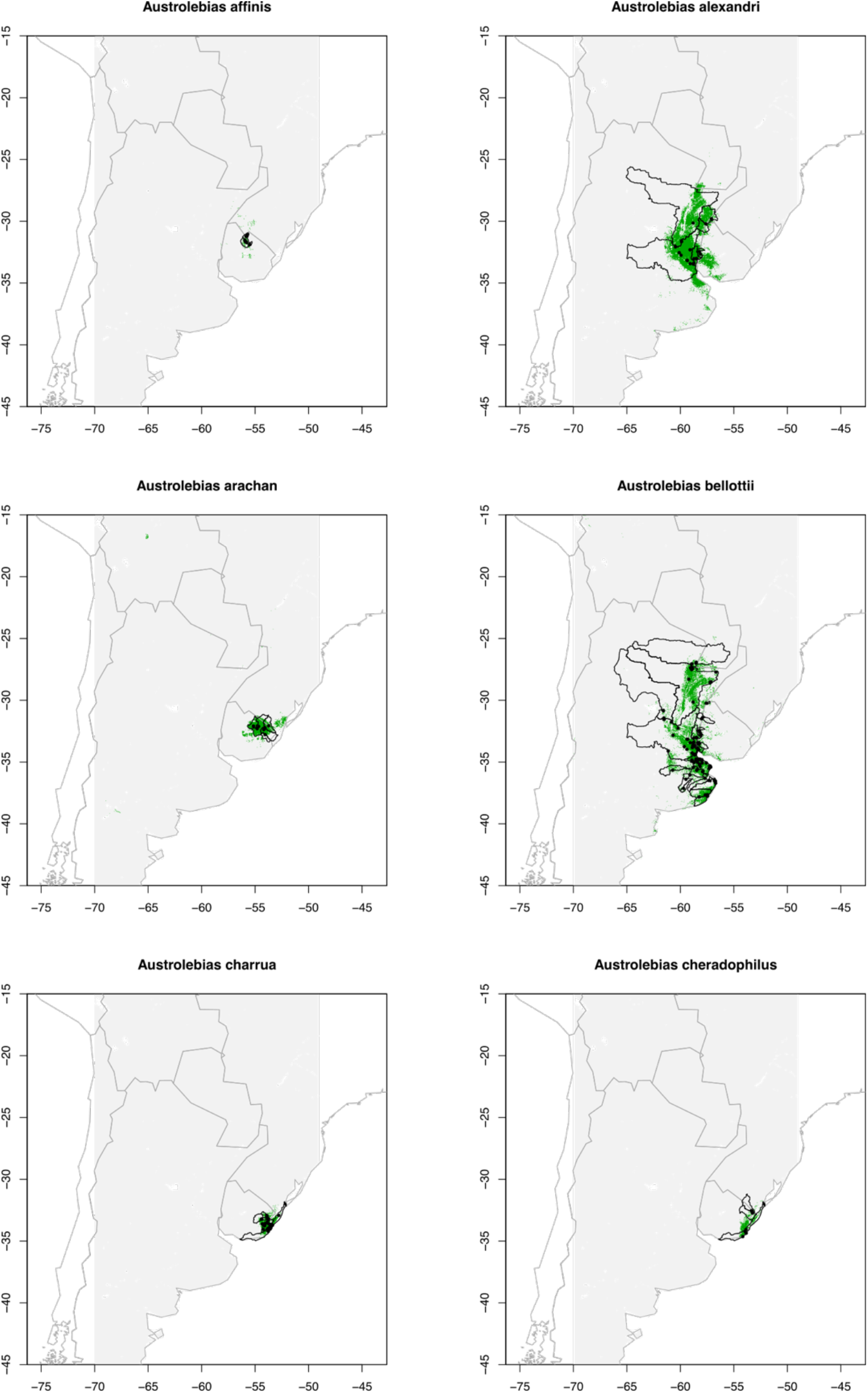
Modelled species ranges. The raster area used in our MaxEnt analyses is shaded grey. Outlines of river basins where species occur in are shown in black (obtained from a global river basin shapefile, with vertices smoothed with a 500m threshold; http://www.waterbase.org/download_data.html. Species occurrences are depicted as black points. Modelled suitability was subjected to a threshold as detailed in the methods section and the resulting predicted ranges are coloured green. The inference for *A. paucisquama* failed.

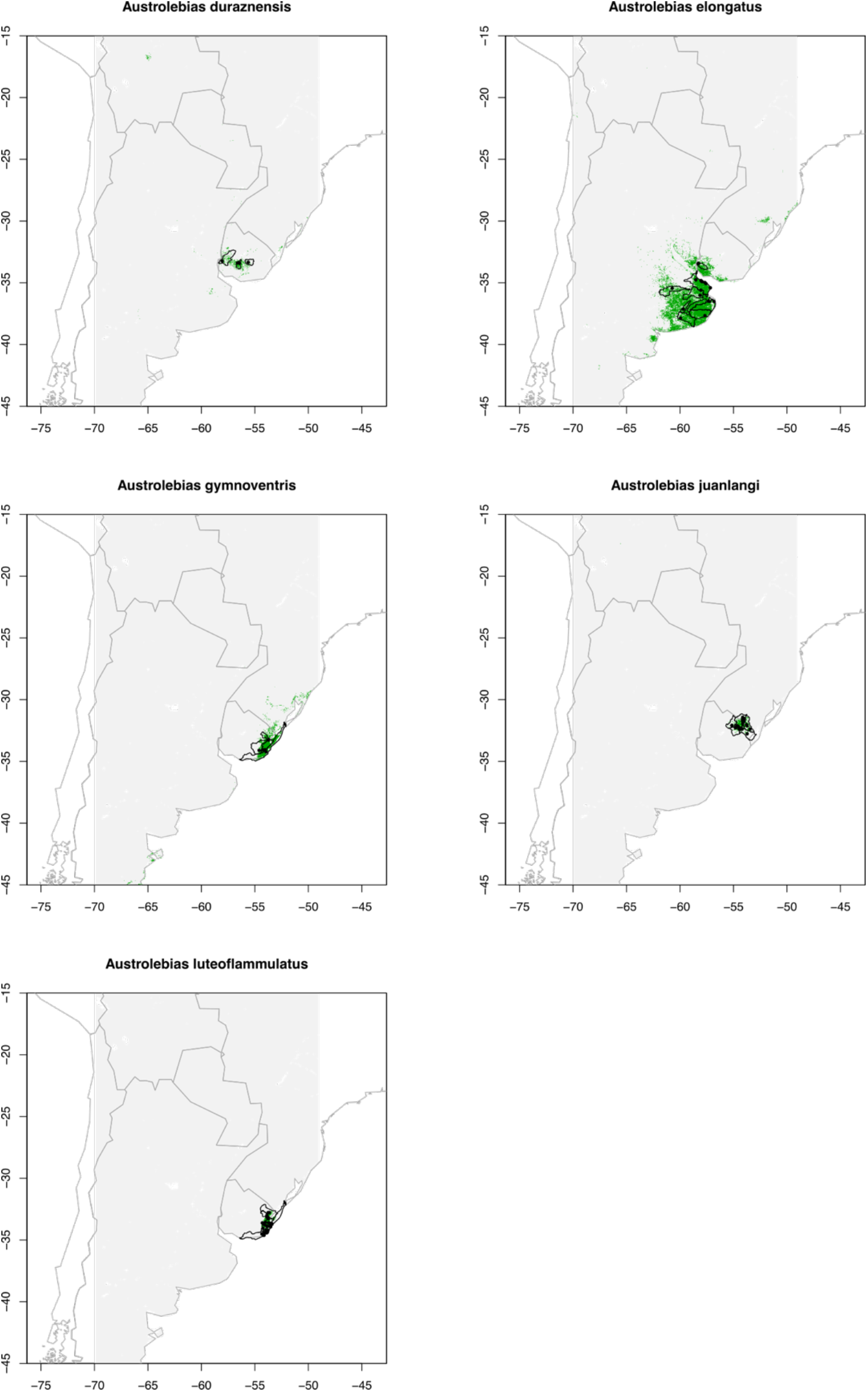

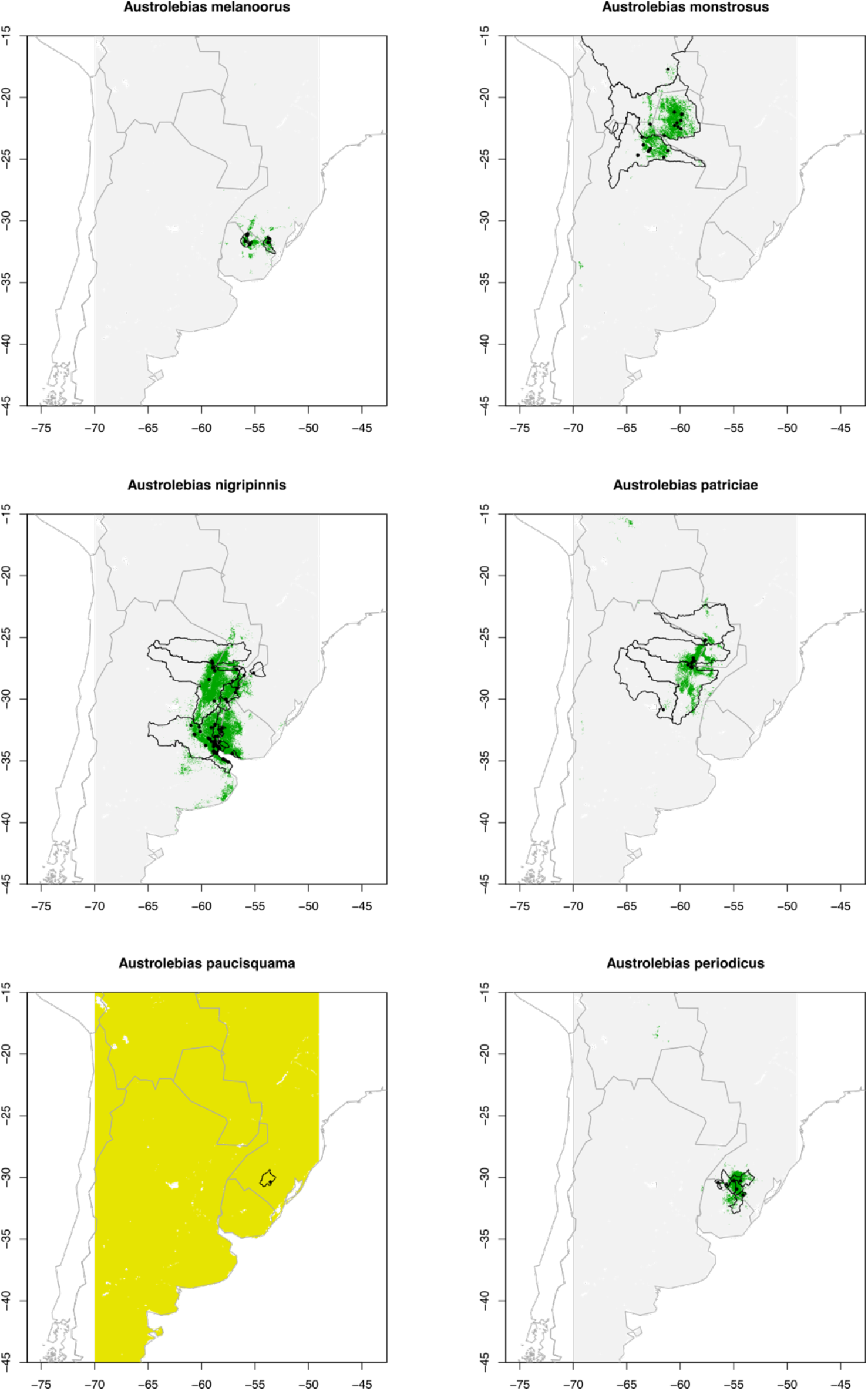

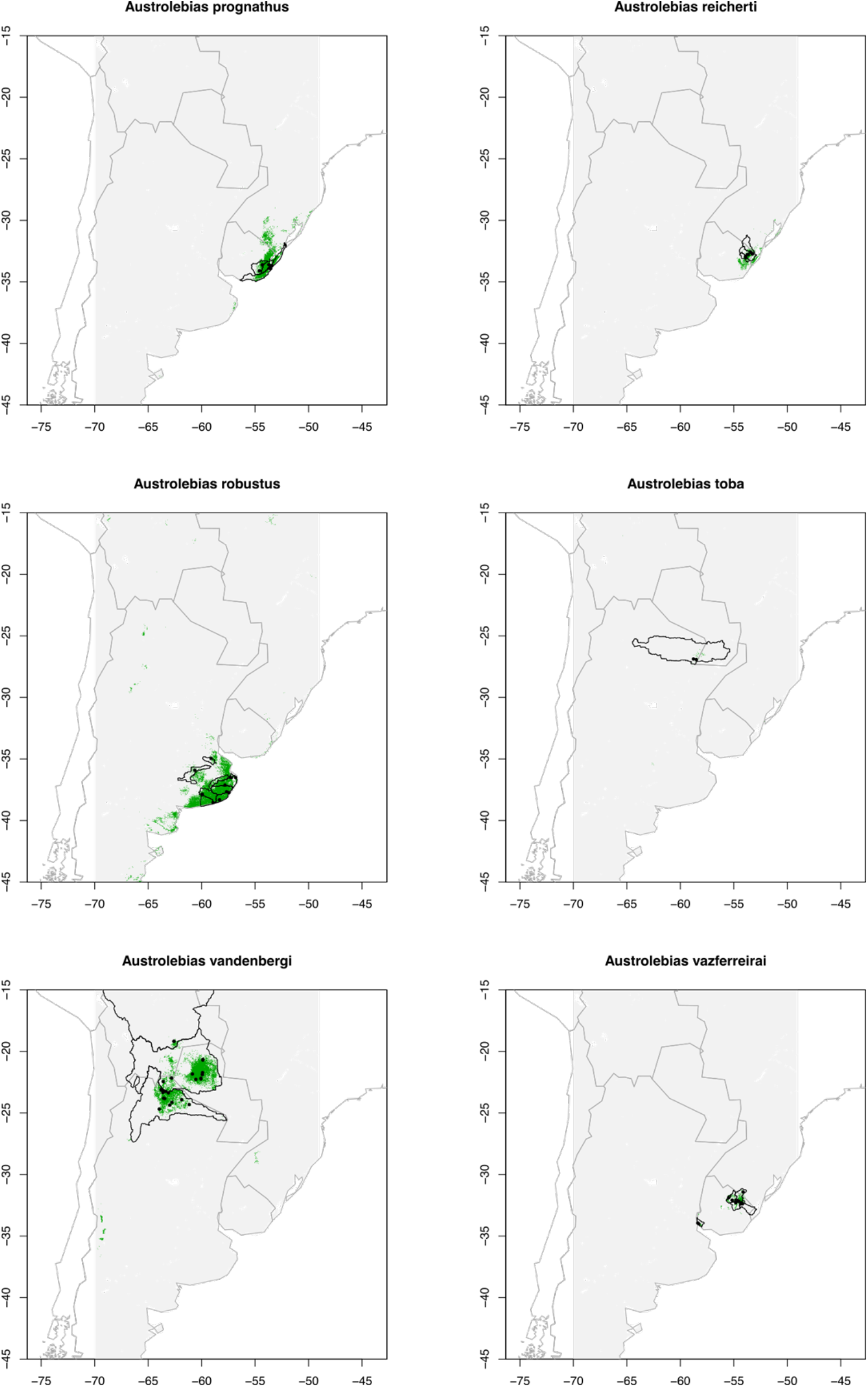

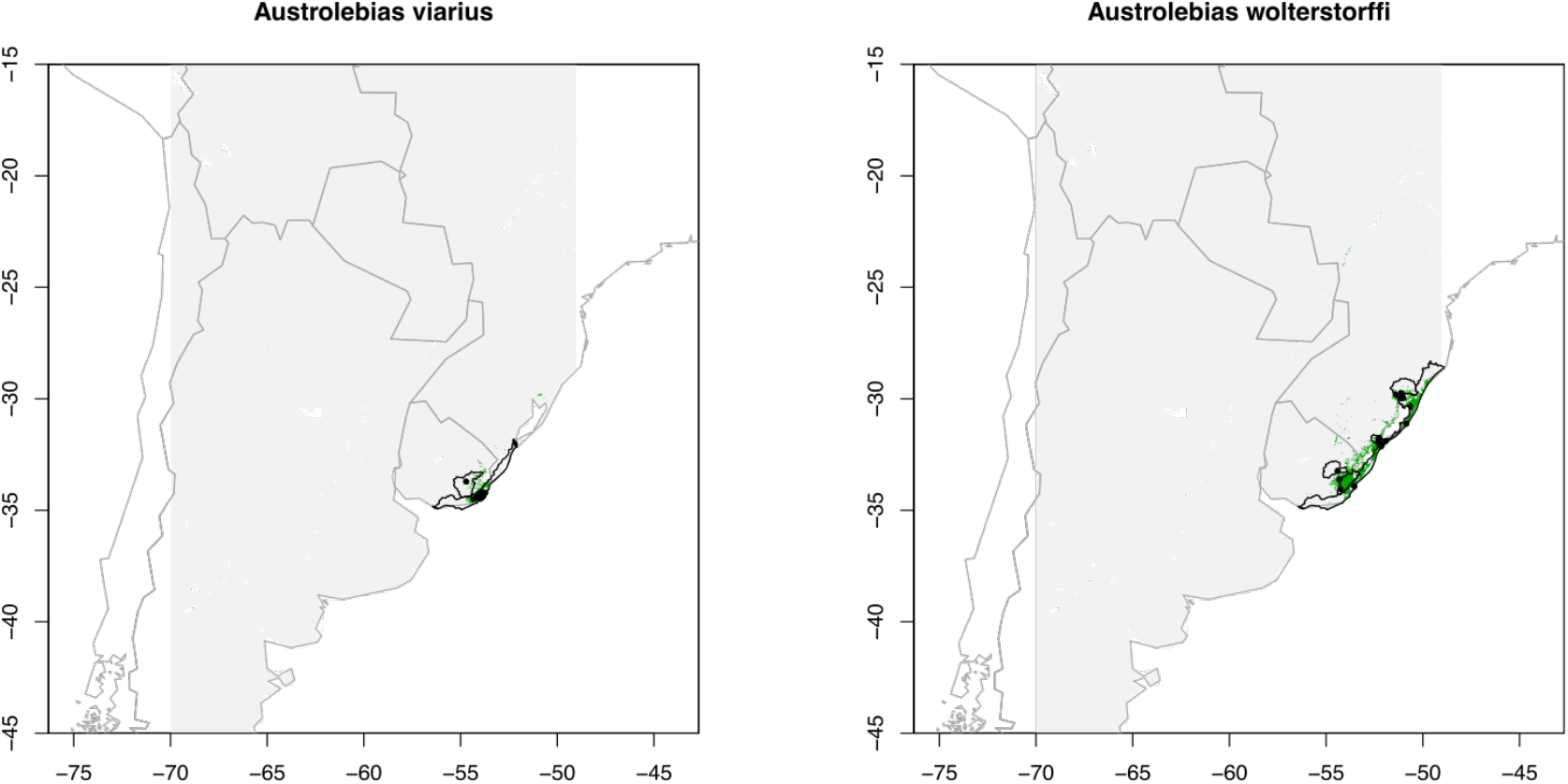

**Figure S7.**
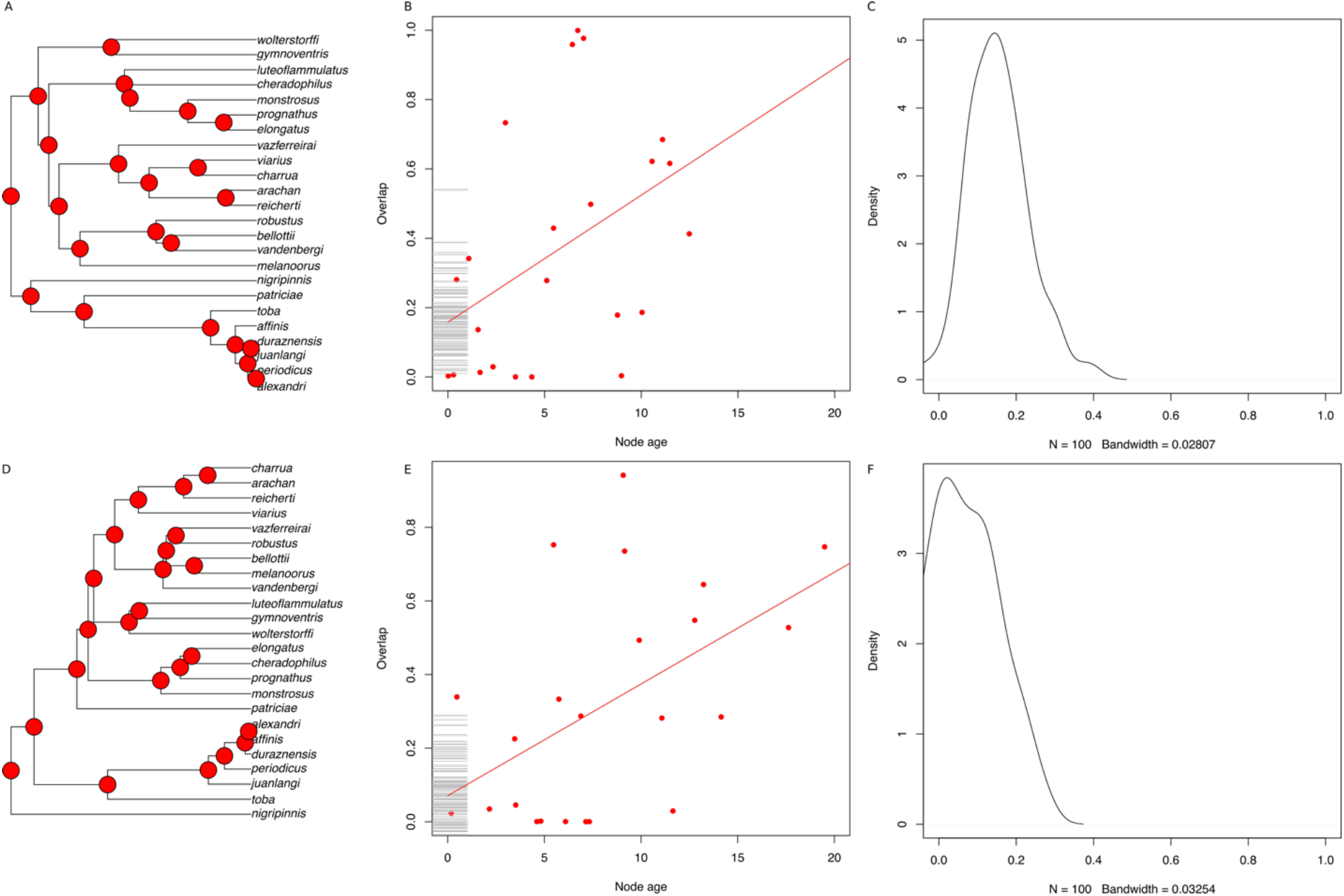
Age-range correlation results depicted for the (A) coalescent tree with pie charts showing predicted cluster membership at each node. Red indicates the cluster with the lowest range overlap, purple intermediate level and blue the highest level of overlap. A scatterplot of node age against range overlap (B) is shown with points coloured by cluster membership. Regression lines fitted to each cluster and bootstrapped intercepts shown as a rug plot along the y axis. A density plot of the inferred intercepts is shown in (C). The same is shown for the mtDNA tree (D), with corresponding scatterplot (E) and density plot (F).

**Figure S8.**
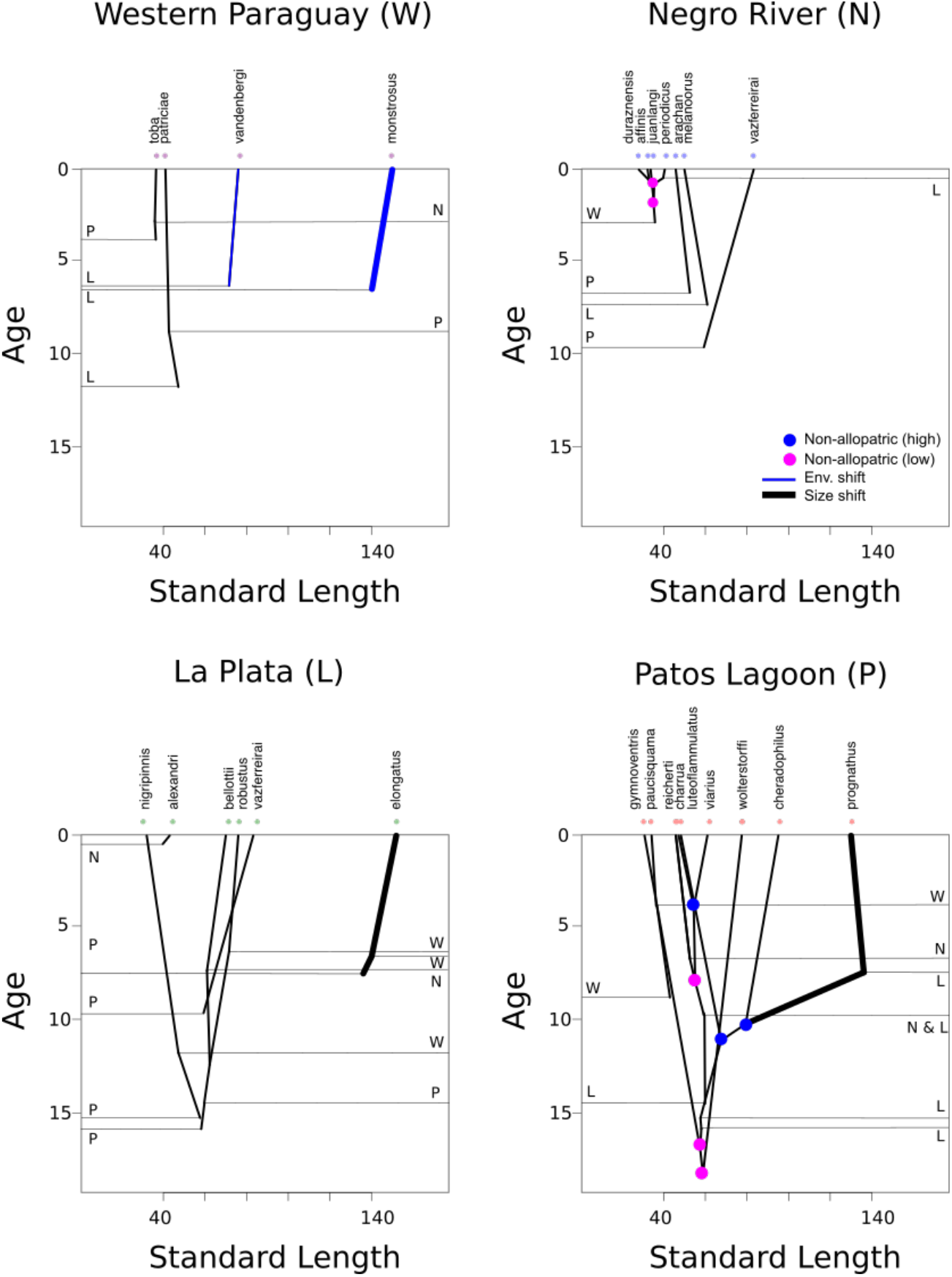
A graphical representation of four species assemblages in *Austrolebias*. Each box represents an area of endemism; age (Ma) is shown on the y axis and maximum standard length (mm) is on the x axis. The nuclear DNA phylogenetic tree is pruned to the species and ancestors that existed in each area based on the results of our ancestral range estimation analysis. Immigration events and source areas are represented on the left of the box and emigrations are shown on the right. Nodes where age-range correlation (ARC) and ancestral range estimation agreed on non-allopatry are coloured based on the cluster they belonged to in our ARC mixture model (Fig. 6A). Shifts estimated by SURFACE are shown blue for environmental niche (see Fig. 5B) and thick branches show the shift in body size optimum (Fig. 5A), or a combination of the two in the case that both shifts were present on the same branch.

**Table S1.**
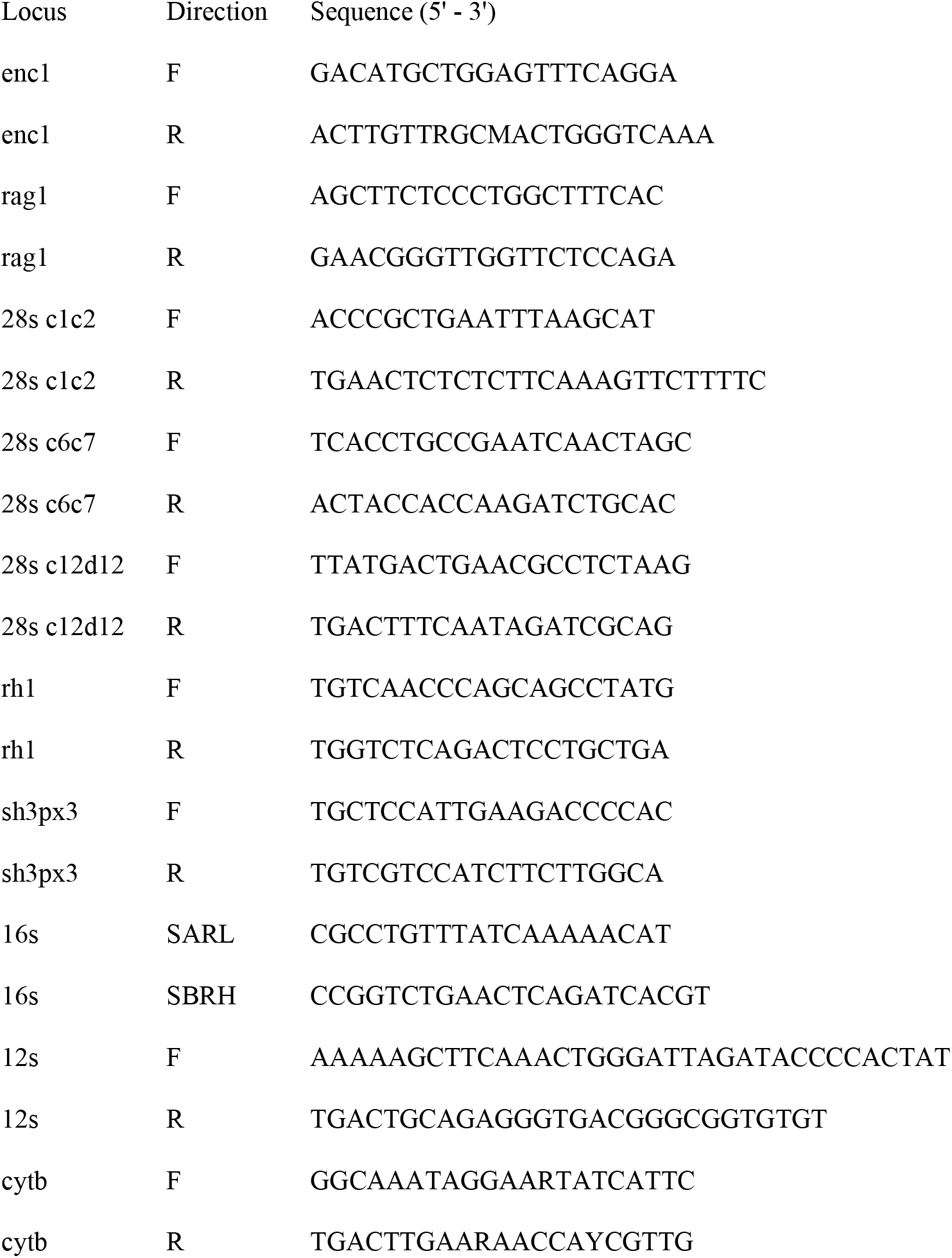
DNA sequences of primers for each locus

**Table S2.**
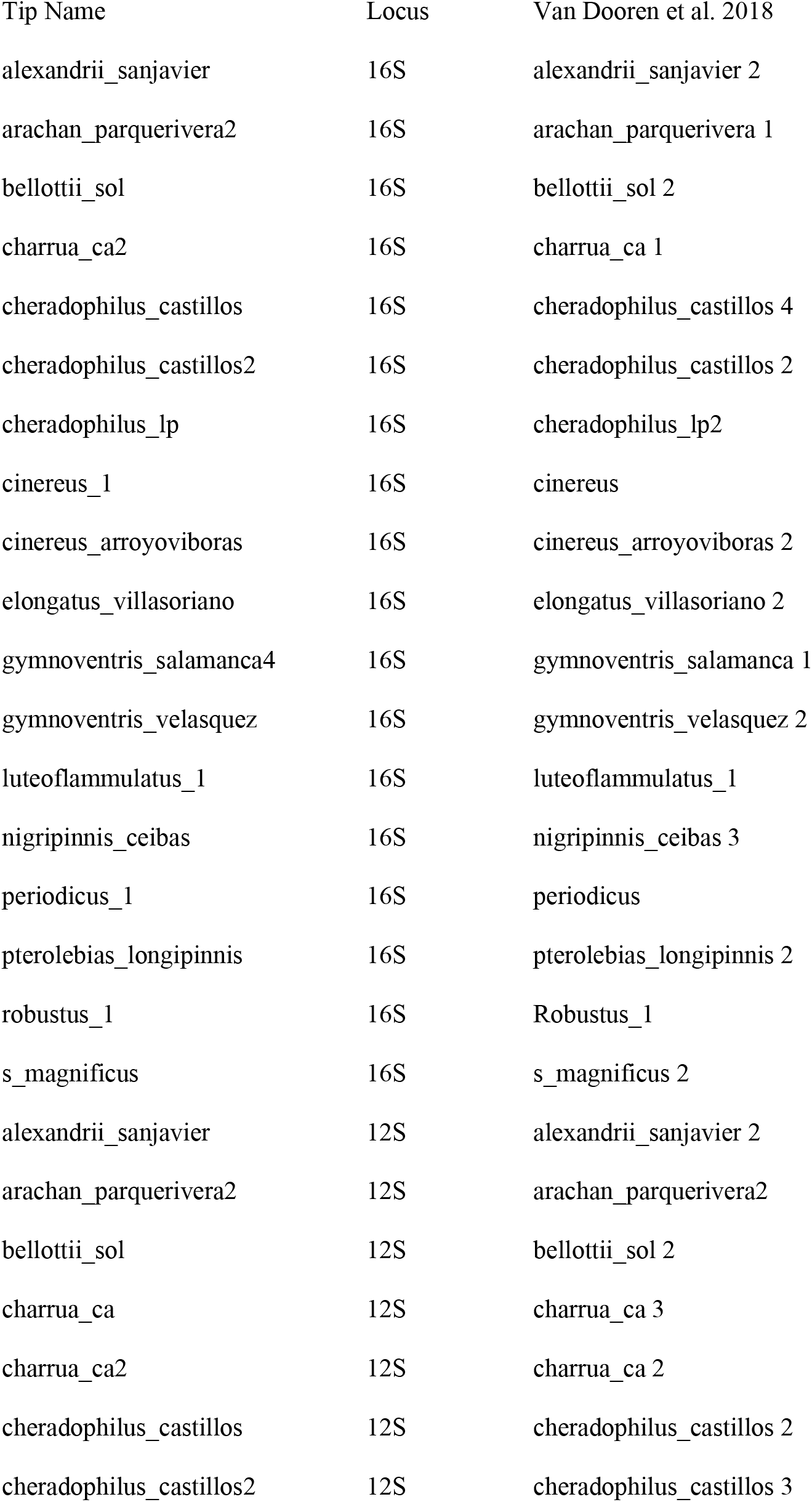
A list of the data taken from Van Dooren et al. (2018) and used for phylogenetic tree inference.

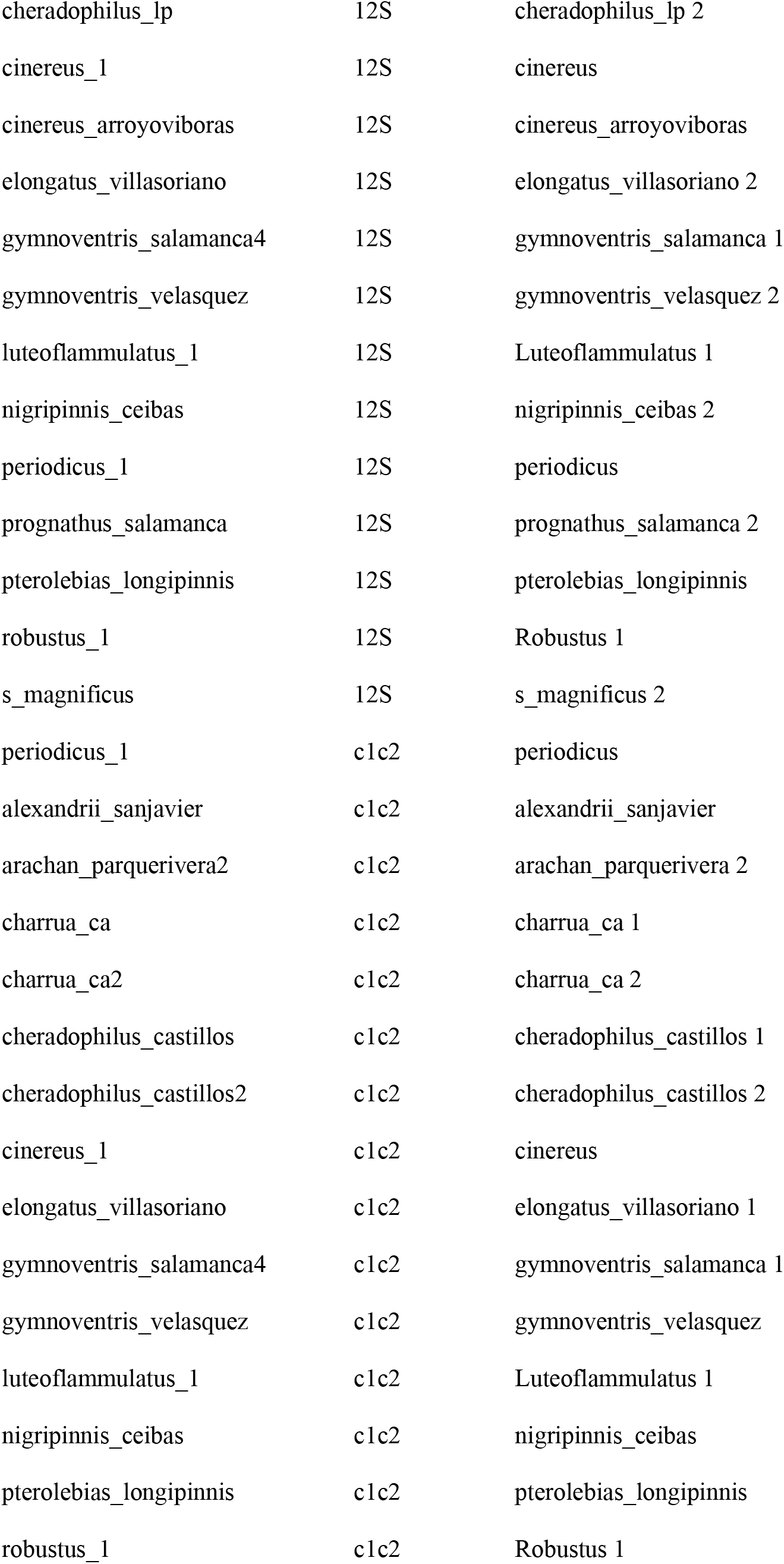

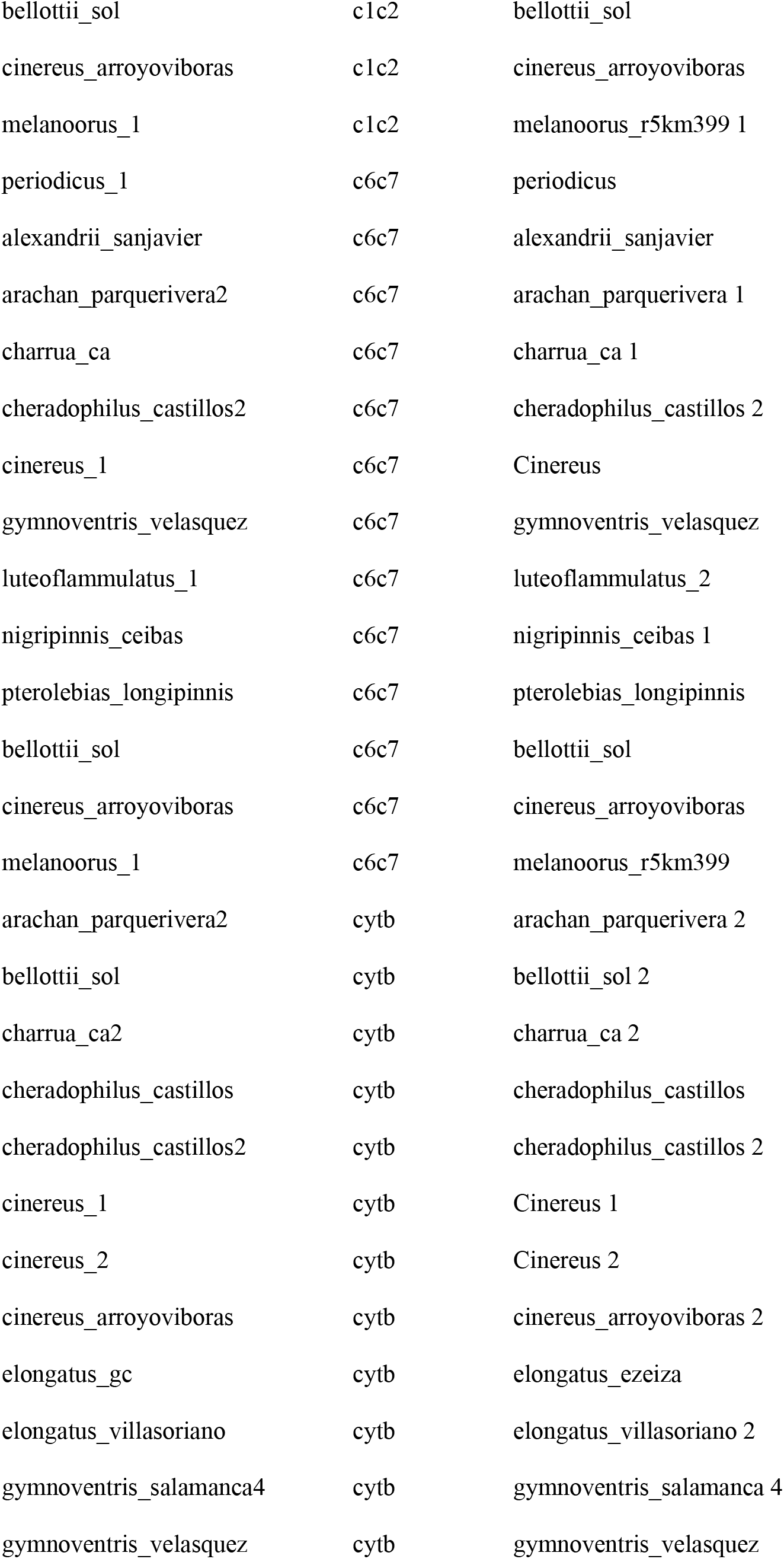

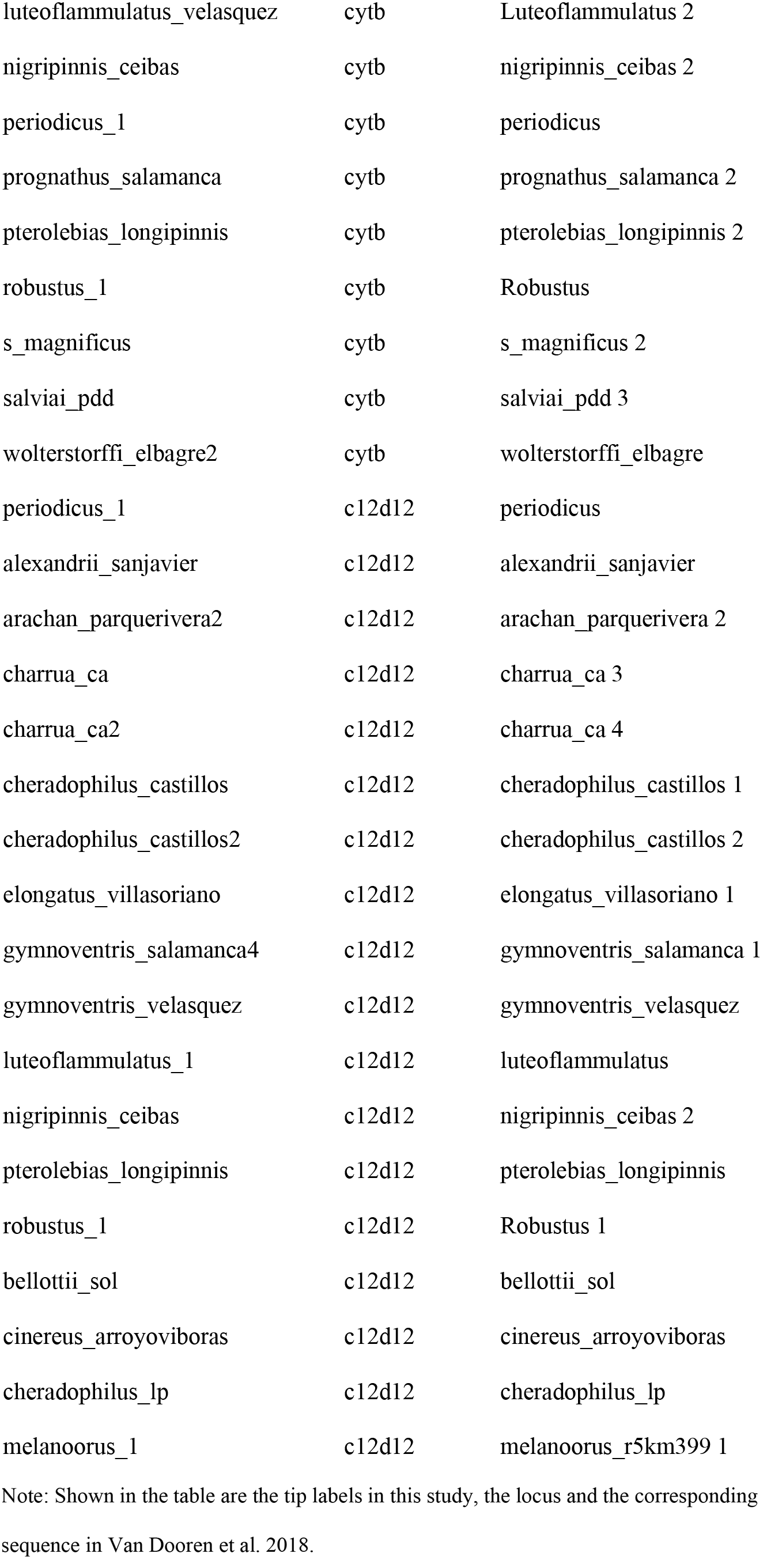

**Table S3** List of *Austrolebias* occurrence locations. Due to the sensitive nature of these data, an excel file of all locations is available upon request.

**Table S4.** Variables used for MaxEnt and PCA analysis of environmental variables at locations where *Austrolebias* have been observed.

| Parameters | Loadings PC1 | Loadings PC2 | Description |
| --- | --- | --- | --- |
| River Basin |  |  |  |
| SLOPE_MEAN | -0.01664 | -0.19114 |  |
| ASPECT_MEAN | -0.15439 | 0.01187 | Mean compass direction slopes face |
| Soil |  |  |  |
| BLDFIE_M_sl2_1km_ll | 0.14784 | 0.06689 | Bulk density (fine earth) in kg / cubic-meter at depth 0.05 m |
| CLYPPT_M_sl2_1km_ll | -0.17428 | -0.10597 | Clay content (0–2 micro meter) mass fraction in % at depth 0.05 m |
| CLYPPT_M_sl6_1km_ll | -0.23710 | 0.02142 | Clay content (0–2 micro meter) mass fraction in % at depth 1.00 m |
| CRFVOL_M_sl2_1km_ll | 0.19497 | 0.09239 | Coarse fragments volumetric in % at depth 0.05 m |
| ORCDRC_M_sl2_1km_ll | -0.12776 | 0.08128 | Soil organic carbon content (fine earth fraction) in g per kg at depth 0.05 m |
| PHIHOX_M_sl2_1km_ll | 0.28586 | -0.04331 | Soil pH x 10 in H2O at depth 0.05 m |
| SLTPPT_M_sl2_1km_ll | 0.18569 | 0.12186 | Silt content (2–50 micro meter) mass fraction in % at depth 0.05 m |
| SLTPPT_M_sl6_1km_ll | 0.23183 | 0.03450 | Silt content (2–50 micro meter) mass fraction in % at depth 1.00 m |
| SNDPPT_M_sl2_1km_ll | 0.02064 | 0.00988 | Sand content (50–2000 micro meter) mass fraction in % at depth 0.05 m |
| SNDPPT_M_sl6_1km_ll | 0.06806 | -0.04879 | Sand content (50–2000 micro meter) mass fraction in % at depth 1.00 m |

Worldclim
|  |  |  |
| --- | --- | --- |
| Altitude | -0.14227 | -0.11809 |
| Annual Mean Temperature | -0.03943 | -0.31807 |
| Mean Diurnal Range | 0.15978 | -0.19825 |
| Isothermality | -0.08156 | -0.24060 |
| Temperature Seasonality | 0.23948 | 0.03849 |
| Max Temperature of Warmest Month | 0.09716 | -0.30108 |
| Min Temperature of Coldest Month | -0.15338 | -0.22317 |
| Temperature Annual Range | 0.23235 | -0.07502 |
| Mean Temperature of Wettest Quarter | 0.09391 | -0.29189 |
| Mean Temperature of Driest Quarter | -0.17140 | 0.00490 |
| Mean Temperature of Warmest Quarter | 0.04840 | -0.31611 |
| Mean Temperature of Coldest Quarter | -0.09713 | -0.29070 |
| Annual Precipitation | -0.27622 | -0.01402 |
| Precipitation of Wettest Month | -0.15459 | -0.21615 |
| Precipitation of Driest Month | -0.25412 | 0.14519 |
| Precipitation Seasonality | 0.16840 | -0.27028 |
| Precipitation of Wettest Quarter | -0.19086 | -0.20828 |
| Precipitation of Driest Quarter | -0.26069 | 0.14317 |
| Precipitation of Warmest Quarter | -0.16828 | -0.22459 |
| Precipitation of Coldest Quarter | -0.25299 | 0.16338 |

Note: Each variable name used is shown as well as a short description in cases where an explanation is needed. Loadings list the contribution of each variable to the scores of the first two principal components used for niche trait analysis.

**Table S5.**
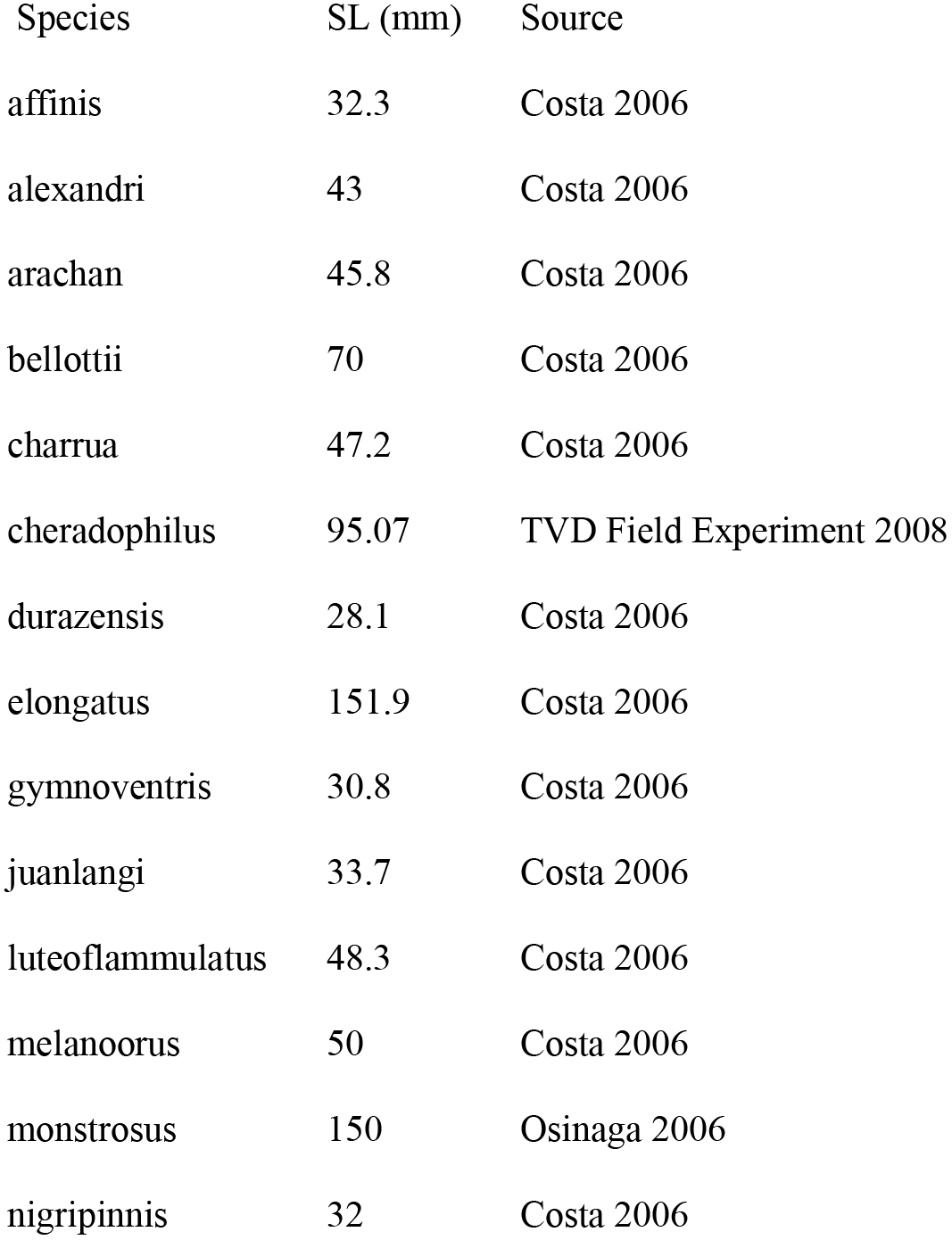
Maximum male body size measurements (standard length) for each species and sources of each measurement.

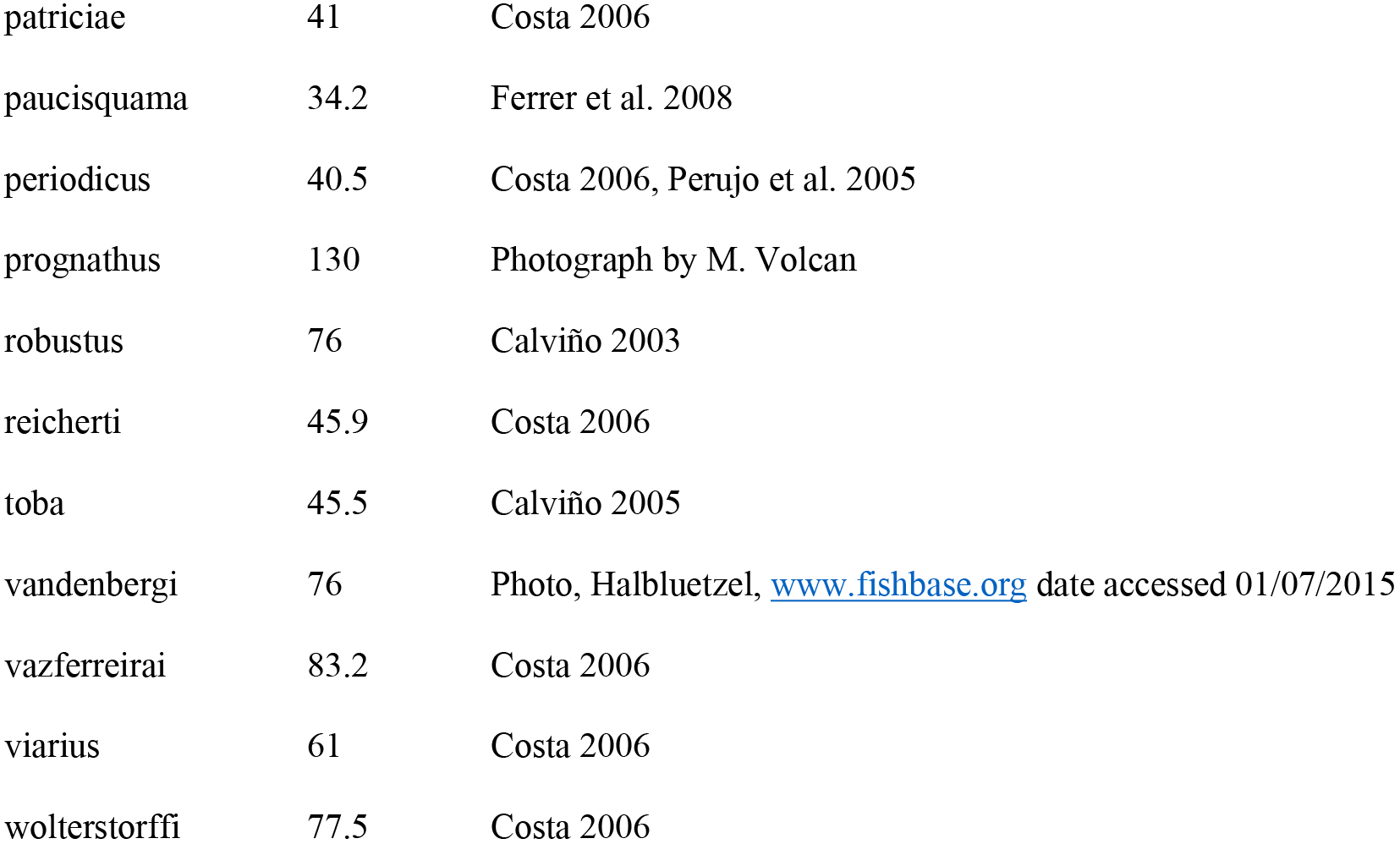

**Table S6.**
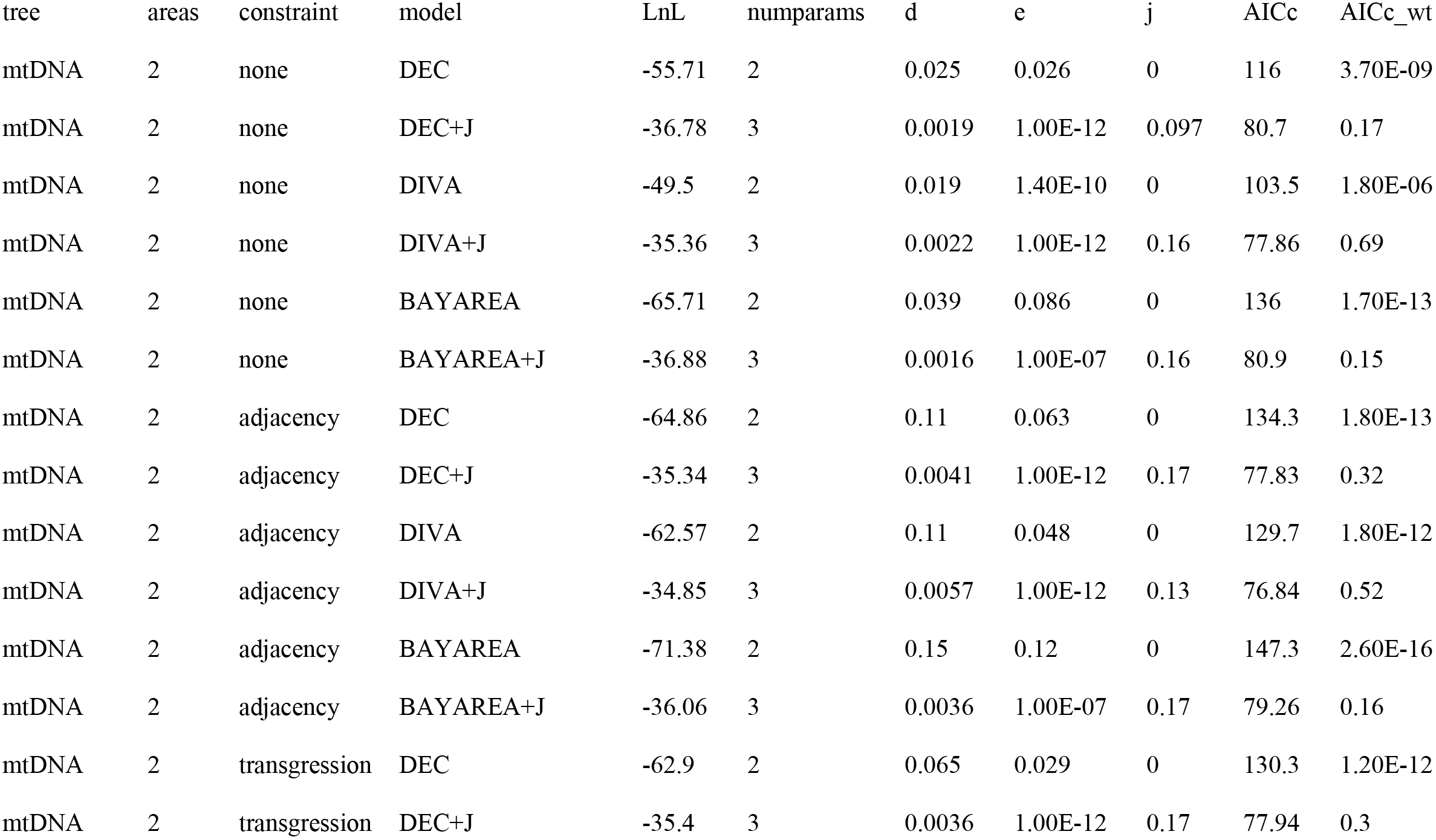
Parameters and statistics from BioGeoBEARS analyses.

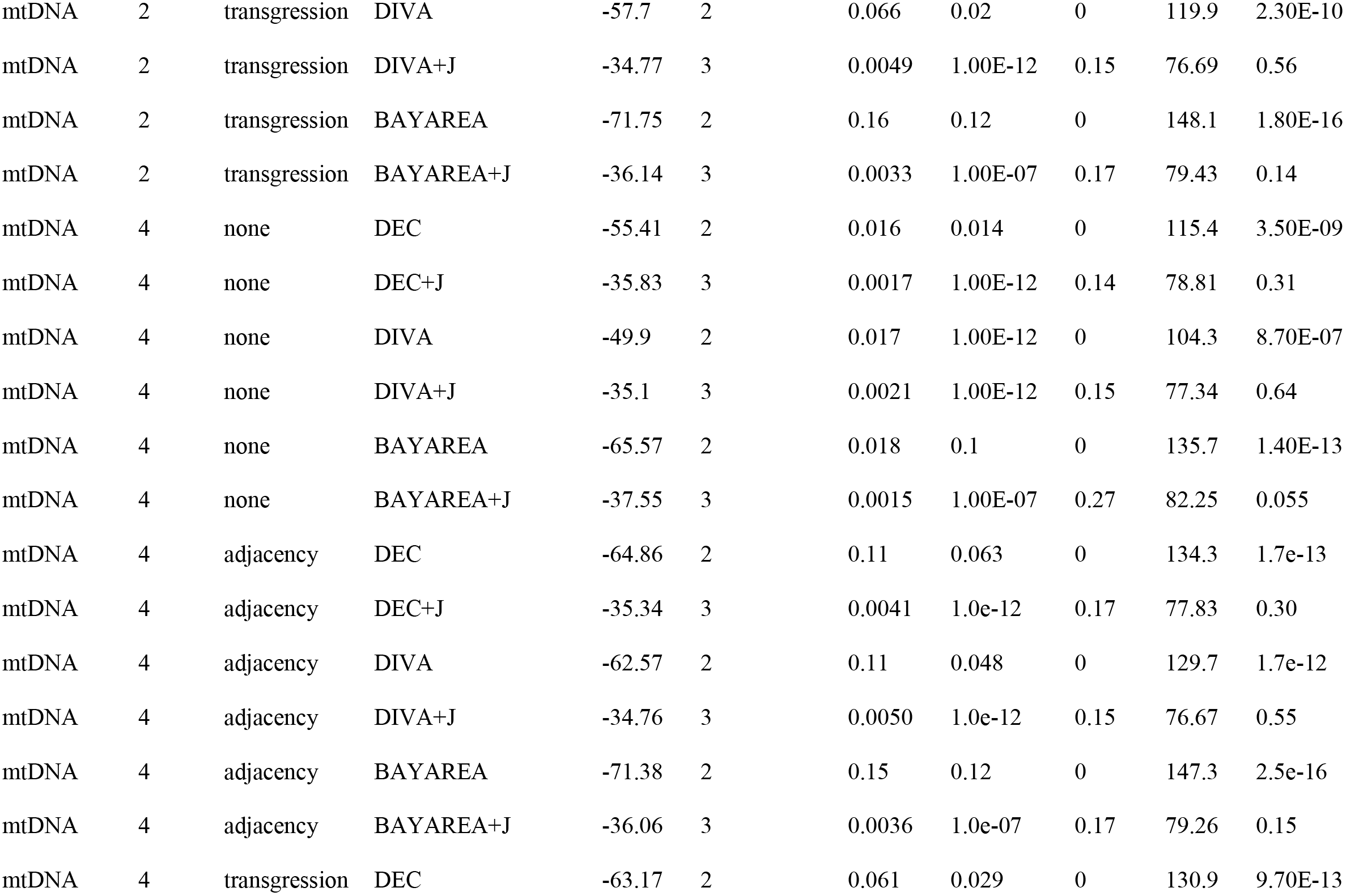

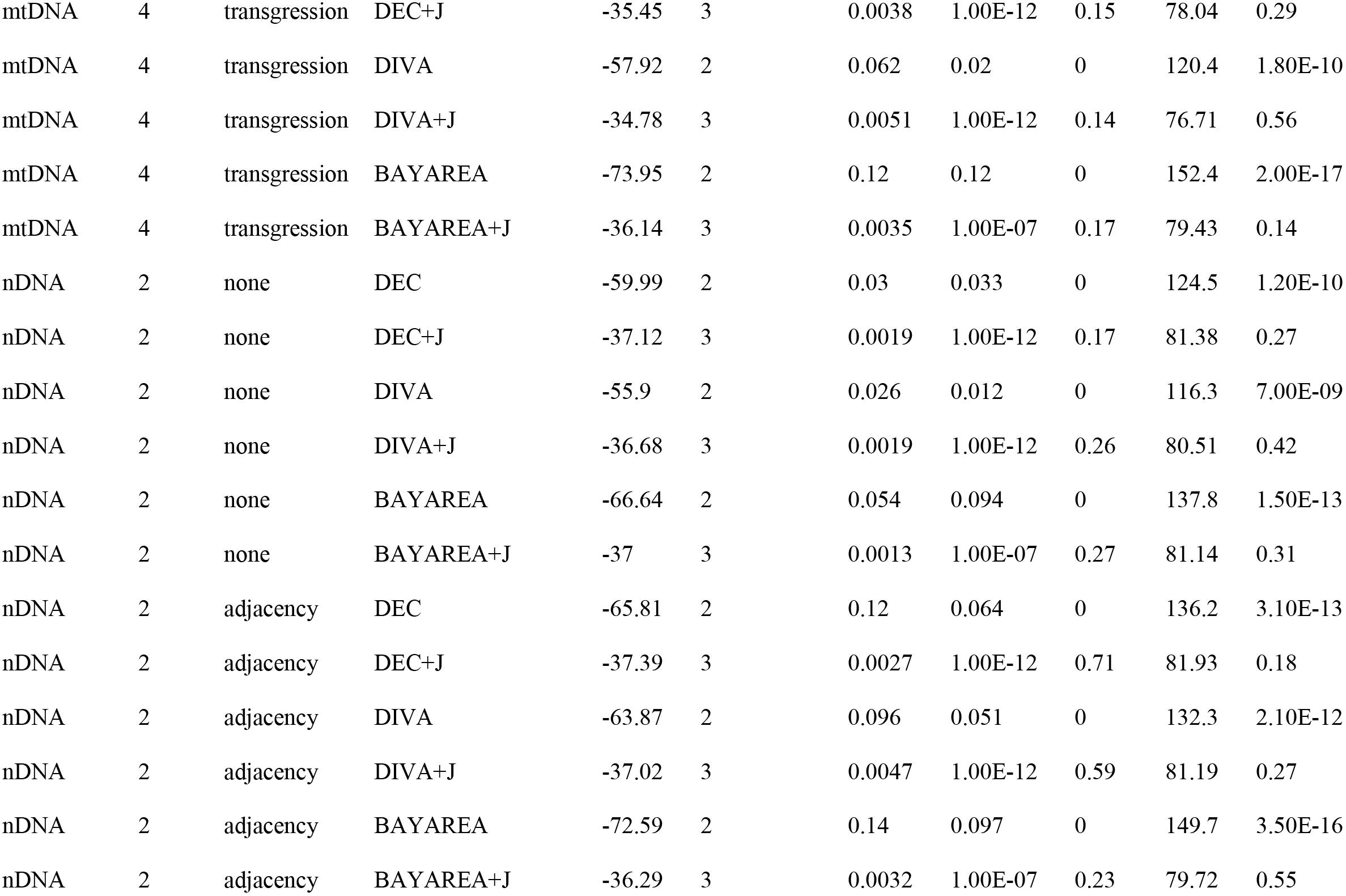

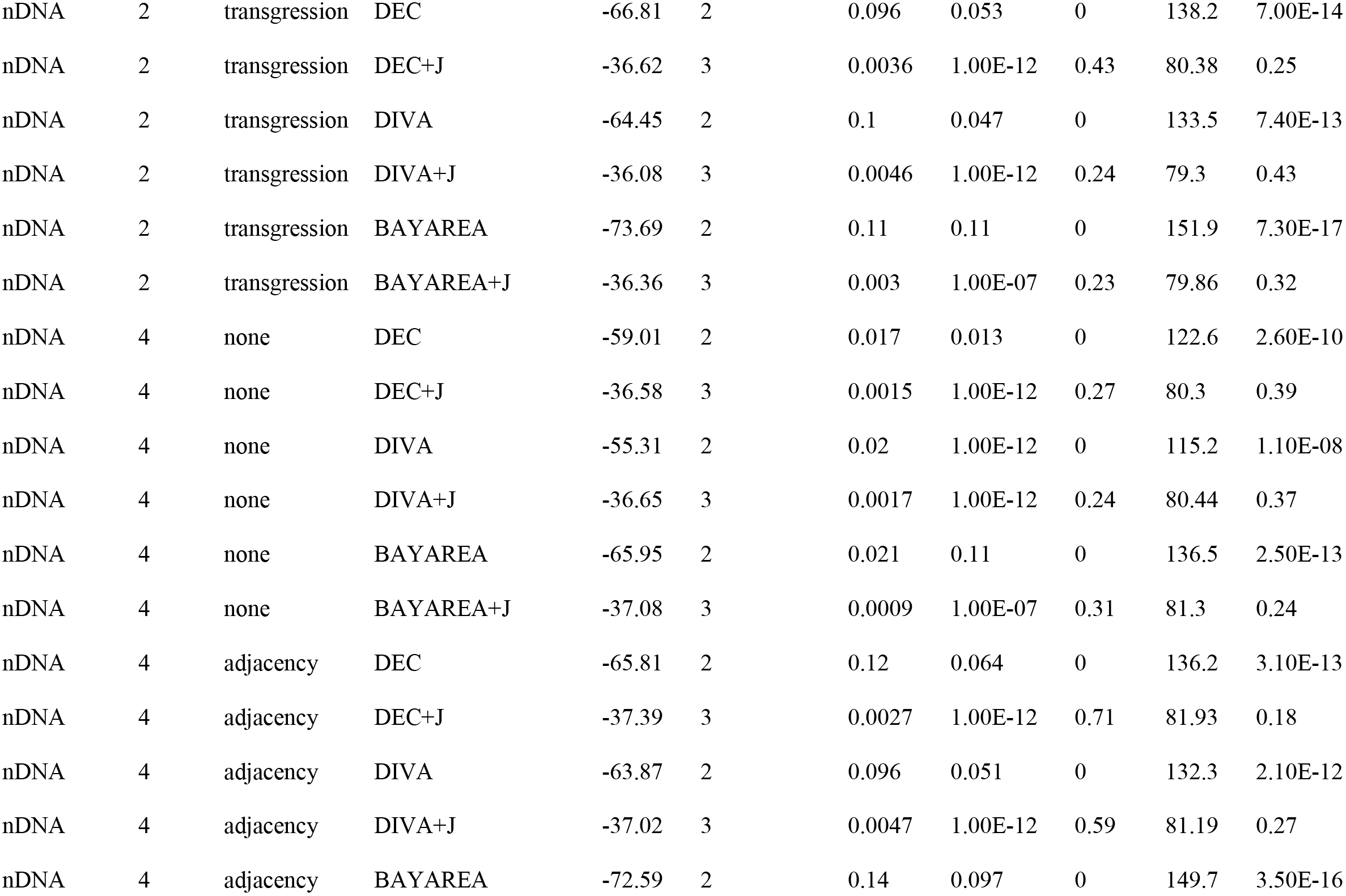

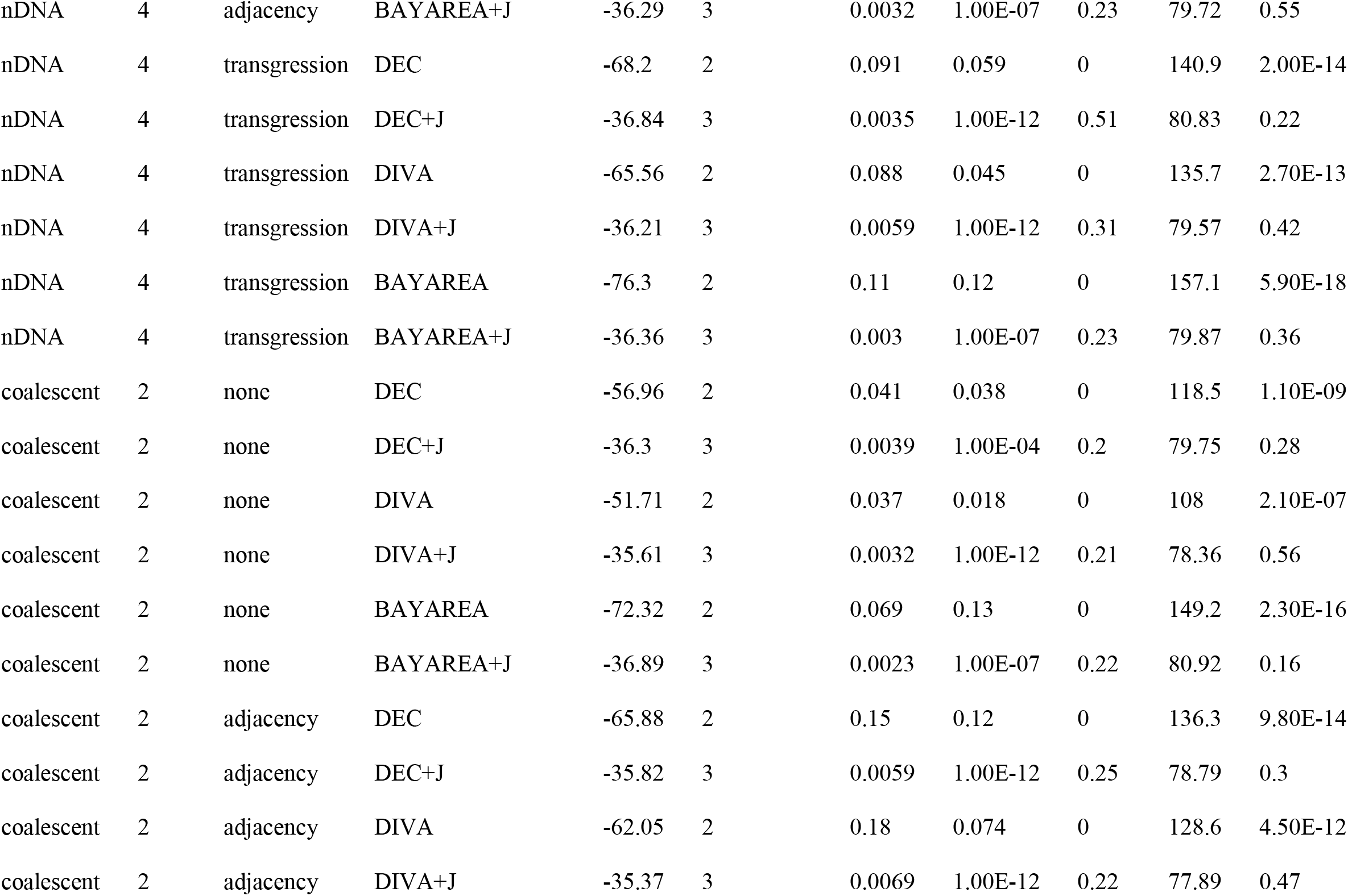

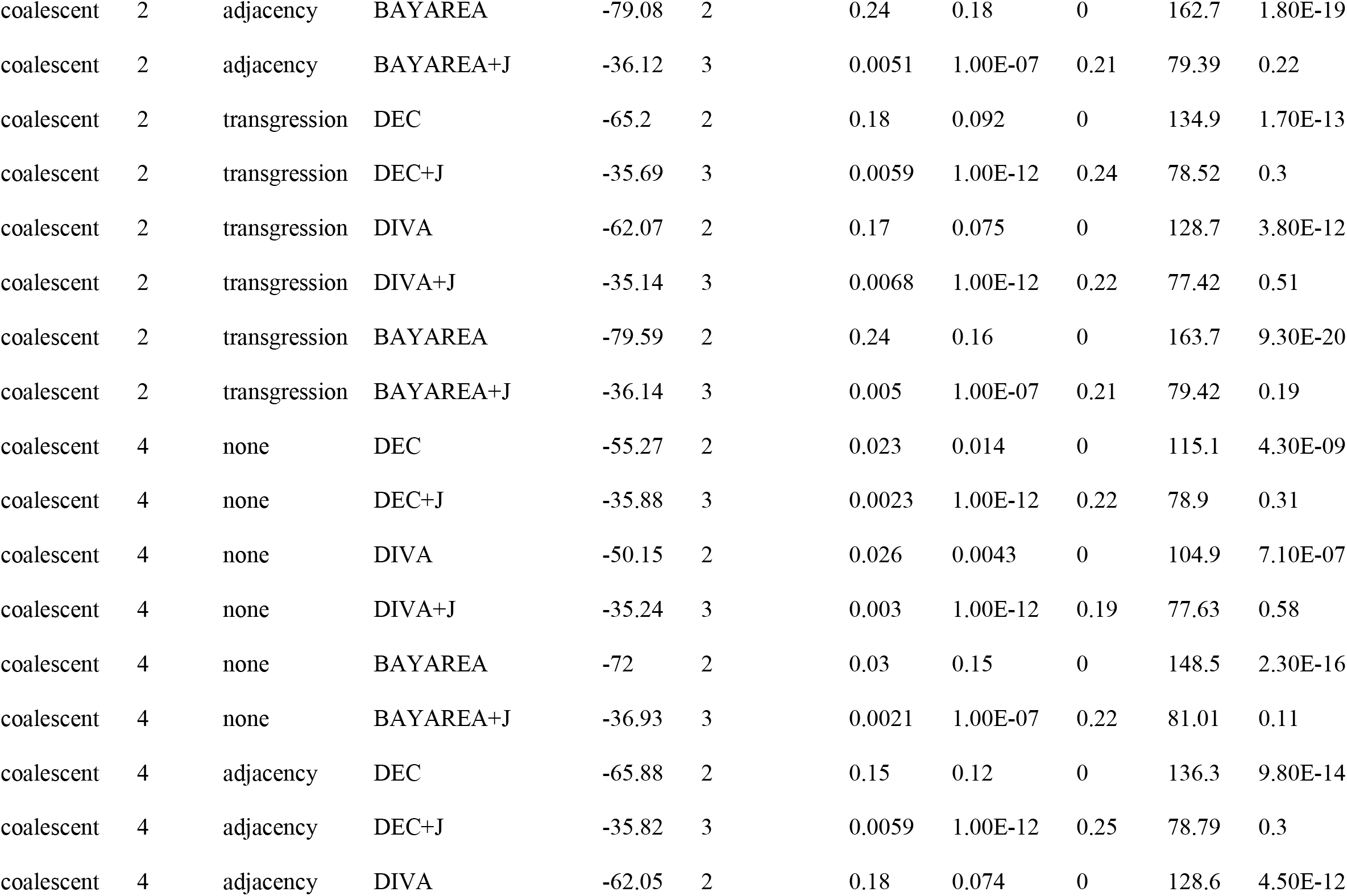

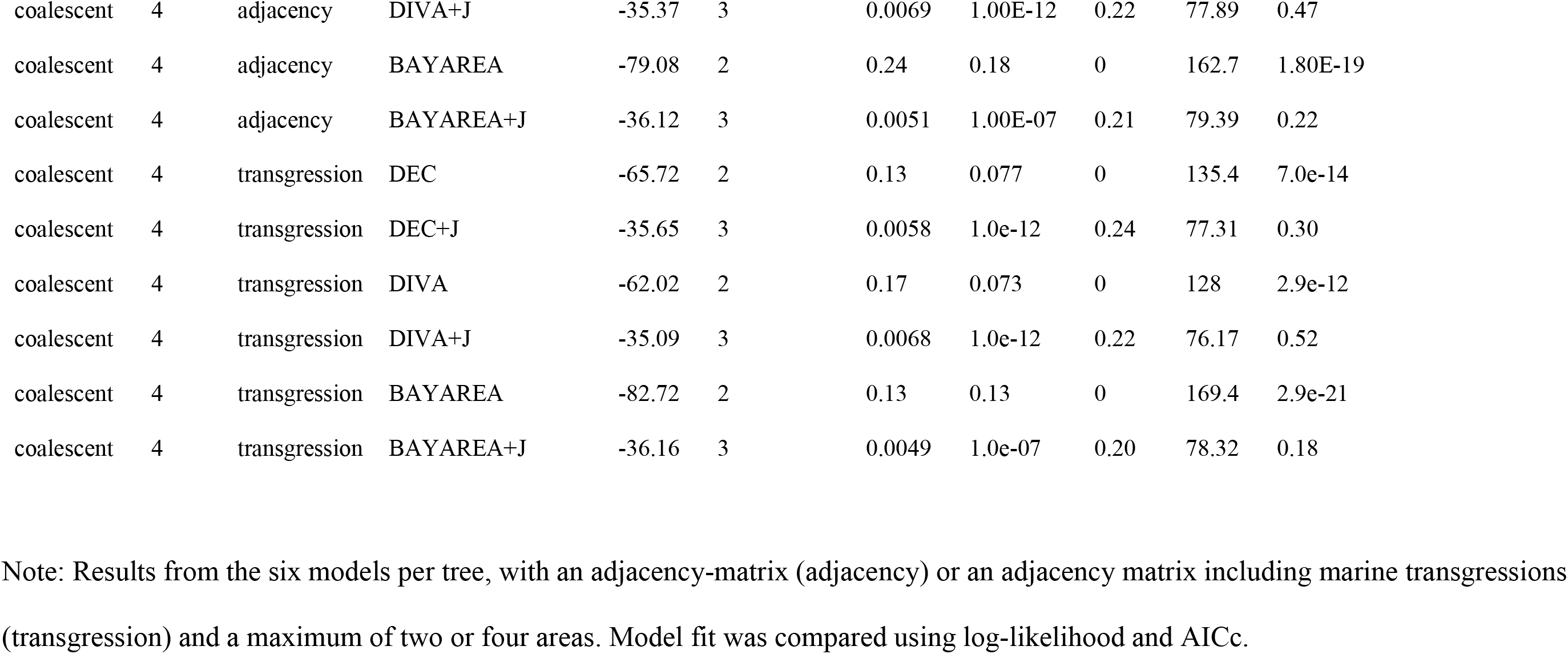

**Table S7.** Comparisons of the best-fitting models with and without the jump-dispersal parameter using likelihood ratio tests.

| data | alt | null | LnLalt | LnLnull | DFalt | DFnull | DF | Dstatistic | pval |
| --- | --- | --- | --- | --- | --- | --- | --- | --- | --- |
| nDNA | DIVA+J | DIVA | -36.68 | -55.9 | 3 | 2 | 1 | 38.43 | 5.70E-10 |
| coalescent | DIVA+J | DIVA | -35.24 | -50.15 | 3 | 2 | 1 | 29.82 | 4.70E-08 |
| mtDNA | DIVA+J | DIVA | -35.36 | -49.5 | 3 | 2 | 1 | 28.28 | 1.00E-07 |

Note: Significant *p*-values indicate that the model with jump dispersal fits better.

**Table S8.**
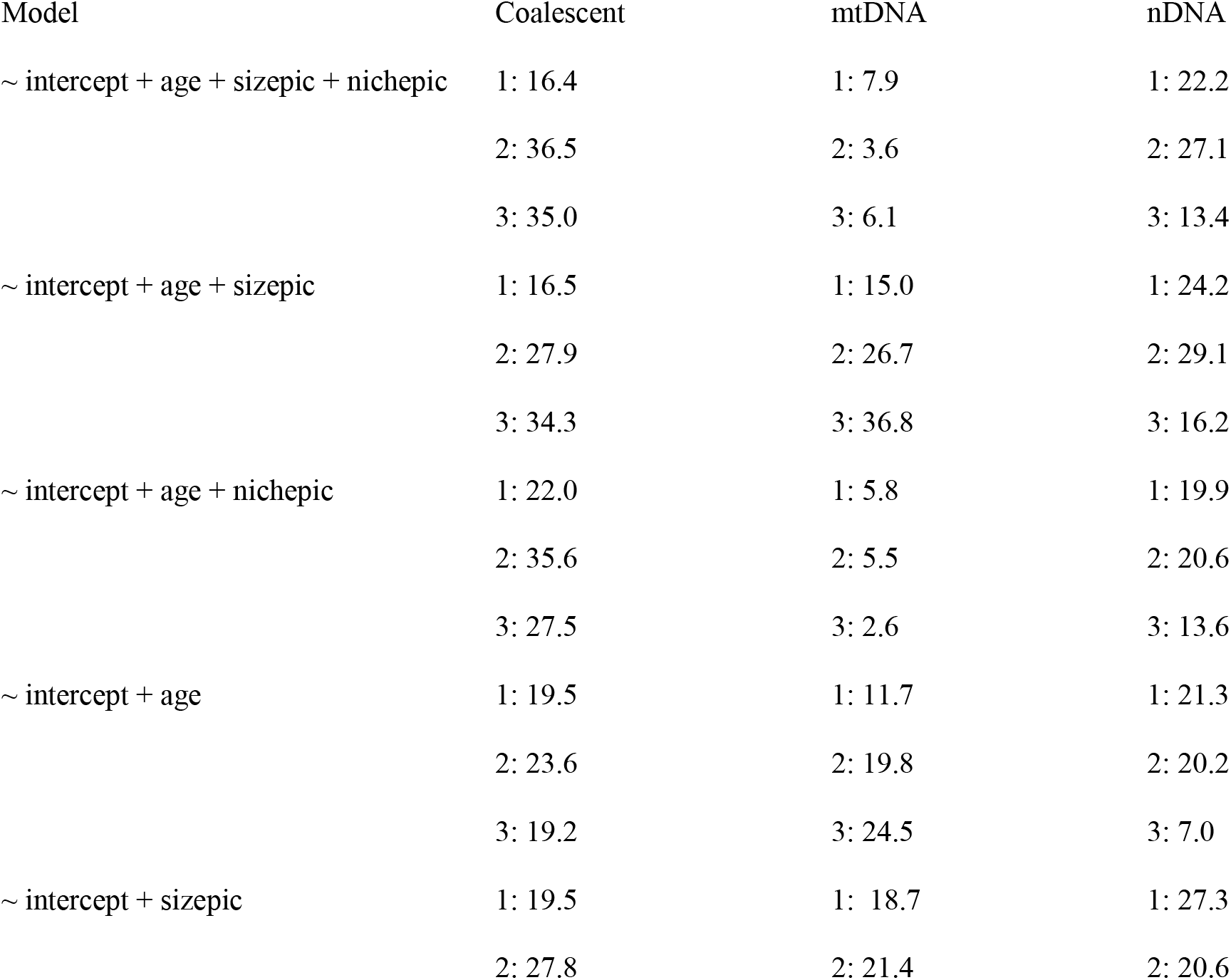
ICL values mixture regressions for ancestral range correlations.

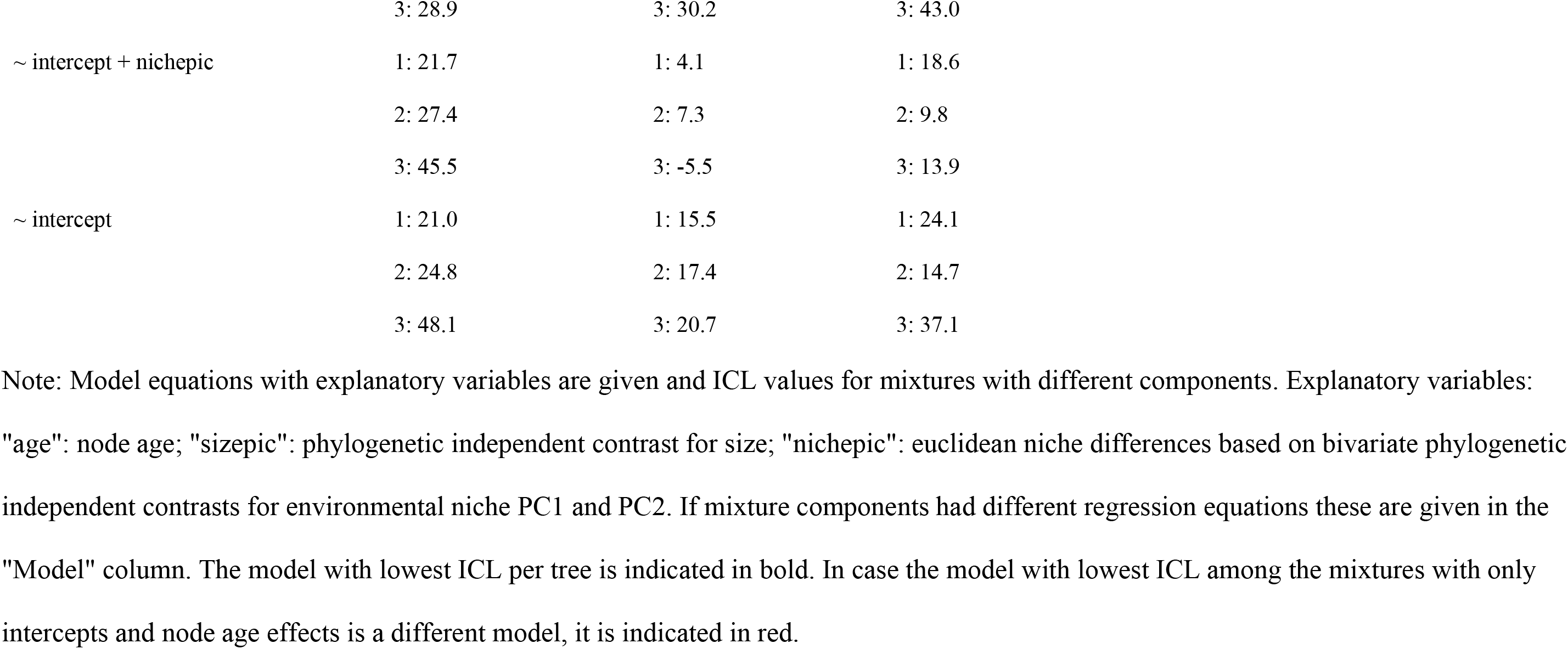

